# A rationally designed and highly versatile epitope tag for nanobody-based purification, detection and manipulation of proteins

**DOI:** 10.1101/640771

**Authors:** Hansjörg Götzke, Markus Kilisch, Markel Martínez-Carranza, Shama Sograte-Idrissi, Abirami Rajavel, Thomas Schlichthaerle, Niklas Engels, Ralf Jungmann, Pål Stenmark, Felipe Opazo, Steffen Frey

## Abstract

Specialized epitope tags are widely used for detecting, manipulating or purifying proteins, but often their versatility is limited. Here, we introduce the ALFA-tag, a novel, rationally designed epitope tag that serves an exceptionally broad spectrum of applications in life sciences while outperforming established tags like the HA, FLAG or myc tags. The ALFA-tag forms a small and stable α-helix that is functional irrespective of its position on the target protein in prokaryotic and eukaryotic hosts. We developed a nanobody (NbALFA) binding ALFA-tagged proteins from native or fixed specimen with low picomolar affinity. It is ideally suited for super-resolution microscopy, immunoprecipitations and Western blotting, and also allows *in-vivo* detection of proteins. By solving the crystal structure of the complex we were able to design a nanobody mutant (NbALFA^PE^) that permits efficient one-step purifications of native ALFA-tagged proteins, complexes and even entire living cells using peptide elution under physiological conditions.

## Introduction

Epitope tags are employed in virtually any aspect of life sciences^1,2^. They are used in biotechnology to facilitate the expression and purification of recombinant proteins^1^ or in cell biology to monitor the biogenesis or spatial organization of a given protein of interest (POI)^3,4^. Other uses include, e.g., the immunoprecipitation of protein complexes studied by mass spectrometry^5,6^, or protein manipulations using tag-binding reagents in living cells^7,8^.

While a given tag might be ideal for a specific application, it may completely fail in others. This is a result of how the tags have been generated – typically as byproducts while developing antibodies against specific POIs (for example the myc-tag^9^, the HA-tag^10^ or the Spot-tag^®11,12^). Other tags, like the His-tag^13^, have been rationally designed for a specific application. None of the available tags covers the full range of current biological applications (see Table S1 for details). For example, the His-tag provides poor results in immunostaining and imaging applications, albeit it is excellent for protein purification. The FLAG, myc and HA-tags have often been used for immunostainings, but due to the large size of the antibodies used as binders, they are suboptimal for super-resolution microscopy and exceedingly difficult to express within cells. These tags therefore cannot be used to reveal or manipulate POIs in living cells. Similar considerations apply for the Twin-Strep tag, which is in addition fixation-sensitive, and therefore not suitable for immunostainings.

More recently the EPEA tag^14^ (also known as C-Tag) and the Spot-tag^®11,12^ have been identified as tags recognized by camelid single-domain antibodies (sdAbs, also known as nanobodies^15^). In contrast to conventional antibodies, sdAbs are monomeric, small, show a superior performance in super-resolution microscopy^16,17^ and can, in principle, even be used for intracellular applications^7,8^. Unfortunately, both of these systems have several problems that override the potential advantages given by their nanobody binders. For instance, both nanobodies detect endogenous proteins (α-synuclein and β-catenin, respectively) and display comparably poor affinities when used as monovalent binders (Table S1) implying that they are suboptimal for advanced microscopy applications or pull-downs of low abundant proteins. Additionally, both tags have so far not been reported to work in living cells, and the EPEA tag can only be used at the C-terminus of target proteins.

To overcome the limitations of the available epitope tags, a first step is to define a set of desired features. A truly versatile tag should not affect the structure, topology, localization, solubility, oligomerization status or polar interactions of the tagged protein^18,19^. It should therefore be small, monomeric, highly soluble^20^ and electroneutral. For highest versatility, it should in addition be resistant to chemical fixation. To avoid specific background signals, the optimal tag should be unique in eukaryotic and prokaryotic hosts, while being well expressed and protease resistant. Similar, the molecule binding such tag should fulfill certain characteristics. It should not only be small for an optimal access to crowded regions and have a minimal linkage error in super-resolution microscopy^21,22^, but also specifically bind the tag with high affinity. For *in-vivo* imaging and manipulations, the binder should be genetically accessible and fold properly within various host organisms. For biochemical applications, the preferred binder should allow for both, stringent washing and native elution of immunoprecipitated proteins or complexes. Strikingly, currently existing epitope tag systems fail to fulfill the complete set of mentioned criteria (Table S1). To manufacture an epitope tag system with the ultimate versatility, the only way is to design it *de novo*.

With this clear objective we created the ALFA-tag. This 15 amino acids sequence is hydrophilic, uncharged at physiological pH and devoid of residues targeted by amine-reactive fixatives or cross-linkers. It has a high propensity to form a stable α-helix that spontaneously refolds even after exposure to harsh chemical treatment. Due to its compact structure, the tag is physically smaller than most linear epitope tags. The ALFA-tag is compatible with protein function and can be placed at the N- or C-terminus of a POI, or even in between two separately folded domains.

As a counterpart binding the ALFA-tag, we developed a high-affinity nanobody (NbALFA), which proved to be suitable for super-resolution imaging and intracellular detection of ALFA-tagged proteins and allowed very efficient and clean immunoprecipitations and Western blots. However, it was virtually impossible to separate NbALFA from the ALFA-tag under native conditions, hindering its application for the purifications of native protein complexes, organelles or cells. Based on the crystal structure of the NbALFA-ALFA peptide complex we engineered a nanobody variant (NbALFA^PE^; “***P***eptide-***E***lution”) suitable for highly specific purification of ALFA-tagged proteins, protein complexes and even living cells under physiological conditions. The rationally designed ALFA system presented here serves an exceptionally broad range of applications from biotechnology to cell biology. A single tag can therefore replace a great variety of traditional epitope tags.

## Results

### The ALFA-system

The ALFA-tag sequence (Fig.1a) is inspired by an artificial peptide reported to form a stable α-helix in solution^23^. It was selected based on the following properties: i) It features a high alpha-helical content, ii) The sequence is absent in common eukaryotic model systems, iii) It is hydrophilic and neutral at physiological pH while retaining moderate hydrophobic surfaces and iv) It does not contain any primary amines that are modified by aldehyde-containing fixatives. In order to develop nanobodies binding the ALFA-tag, we immunized alpacas and selected nanobodies specifically targeting ALFA-tagged proteins by a novel nanobody selection method (see Online Methods). The best binder (NbALFA; Fig.1b) was genetically modified with ectopic cysteines allowing for a site-specific immobilization or fluorophore attachment^24^.

**Fig. 1.**
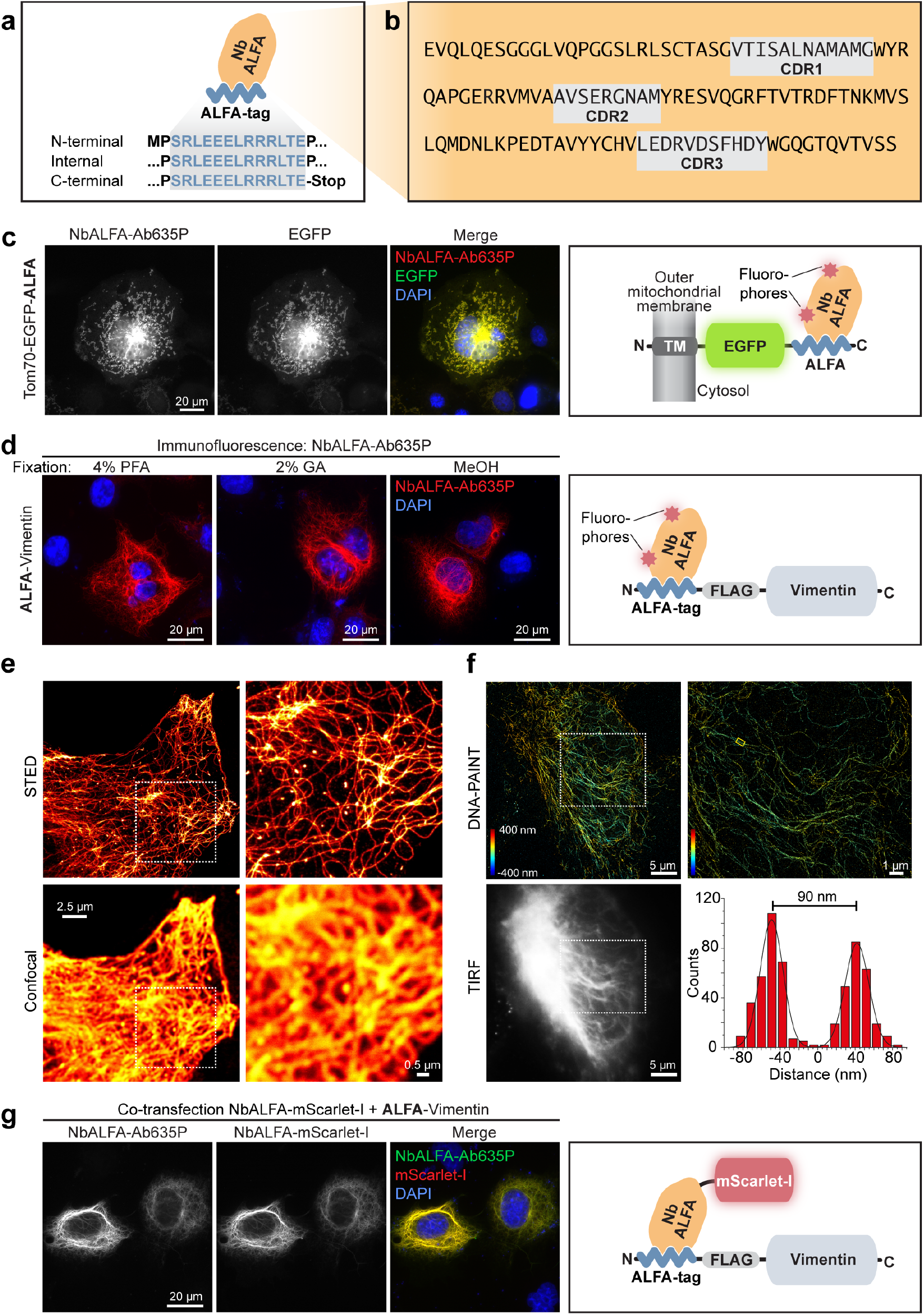
Nanobody-based detection of ALFA-tagged proteins in immunofluorescence applications. **a;** Sketch of NbALFA bound to ALFA-tags with ALFA sequences for tagging at various positions. **b;** Sequence of NbALFA. Grey boxes indicate CDRs 1-3 (AbM definition). **c;** COS-7 cells transfected with Tom70-EGFP-ALFA were fixed with paraformaldehyde (PFA) and stained with NbALFA coupled to AbberiorStar635P (NbALFA-Ab635P). Left to right: NbALFA-Ab635P; intrinsic EGFP signal; overlay incl. DAPI stain; sketch illustrating the detection of Tom70-EGFP-ALFA. **d**; ALFA-Vimentin was detected with NbALFA-Ab635P after fixation with 4% PFA, 2% glutaraldehyde (GA), or 100% Methanol (MeOH). **e**, STED and confocal images of COS-7 cell transiently transfected with ALFA-Vimentin and stained with NbALFA-Ab635P. **f;** HeLa cells transfected with ALFA-vimentin were stained with NbALFA bearing a 10-nucleotide single stranded DNA before imaging by 3D DNA-PAINT. The histogram refers to a region (small yellow rectangle) where 2 vimentin filaments are resolved although being only ~90nm apart. The localization precision was 5.2nm. **g;** COS-7 cells were co-transfected with an NbALFA-mScarlet-I fusion and ALFA-Vimentin. NbALFA-mScarlet-I co-localizes with ALFA-Vimentin detected by immunofluorescence using NbALFA-Ab635P. This shows that NbALFA expressed in the cytoplasm of mammalian cells can be used for targeting ALFA-tagged proteins in living cells.

### Detection of ALFA-tagged proteins by direct immunofluorescence

In immunofluorescence (IF) applications on PFA-fixed mammalian cells, fluorescently labeled NbALFA specifically recognized target proteins harboring ALFA tags at different locations within the proteins (Fig.S1). Importantly, all tested proteins showed their characteristic localization (Tom70-EGFP-ALFA: mitochondrial outer membrane; ALFA-Vimentin: filamentous structures; EGFP-ALFA-TM: plasma membrane), the ALFA-tag was therefore compatible with the folding and proper localization of the tagged proteins. In a quantitative assay, the nucleocytoplasmic distribution of EGFP carrying an ALFA tag at either terminus was indistinguishable from non-tagged EGFP (Fig.S2). An atypical interaction to cellular membranes or organelles was not observed. Moreover, gel filtration of a recombinant ALFA-tagged mEGFP variant confirmed its monomeric state indicating that the ALFA-tag does not induce multimerization (data not shown). We therefore concluded that the ALFA-tag seems not to impair the physiological behavior of the tagged POIs.

### Resistance to amine-reactive fixatives

In order to facilitate optimization of fixation conditions, it is advantageous if a given epitope tag is compatible with various fixation procedures. In contrast to many established epitope tags (Table S1), the ALFA-tag does not contain lysines by design. Consequently, it could be detected after standard fixation with paraformaldehyde or methanol, and was even resistant to 2% glutaraldehyde (Fig.1d). The ALFA-tags is thus compatible with standard fixation methods and has the potential to be employed in electron microscopy, where glutaraldehyde is preferred due to its superior preservation of cellular structures.

### Super-resolution microscopy

Due to the small size of both the ALFA-tag and NbALFA, the ALFA system results in a minimal linkage-error and it thus perfectly suited for super-resolution microscopy. As examples, we imaged cells transfected with ALFA-Vimentin by either STED microscopy^16^ or 3D DNA-PAINT^25^ using NbALFA directly coupled to a “STEDable” fluorophore or a short single-stranded oligonucleotide, respectively (Fig.1e,f). Our results show that the ALFA system is compatible with the demanding conditions of these fluorescent super-resolution microscopy techniques.

### Detecting and manipulating ALFA-tagged proteins *in vivo*

Some nanobodies are functional within mammalian cells and can thus be used to detect or manipulate target proteins in living cells^8,26^. Indeed, when co-expressing ALFA-tagged target proteins and NbALFA fused to mScarlet-I^27^ in mammalian cells, NbALFA-mScarlet-I robustly co-localized with ALFA-tagged target proteins with minimal off-target signals, resulting in crisp detection of ALFA-tagged structures (Fig.1g, Fig.S3). This demonstrates the ability of NbALFA to bind ALFA tagged proteins in living cells. This feature is very attractive to be used in combination with genome editing tools like CRISPR-Cas^28^ and allows manipulation of ALFA-tagged proteins *in vivo* at their endogenous levels.

### Western Blot

We next tested the ALFA system in Western blots applications and could specifically detect ALFA-tagged Vimentin in lysates from transfected cells using NbALFA labeled with IRDye800CW (Fig.2a, Fig.S4a). To compare the performance of the ALFA system with common epitope tags, we employed a fusion protein harboring HA, myc, FLAG^®^ and ALFA tags (Fig.2b). At identical concentrations of primary antibody or nanobody, the obtained ALFA-tag signal was 3-10-fold stronger than the signal obtained for all other epitope tags (Fig.2c, Fig.S4b). This is striking since the ALFA-tag signal exclusively relied on the NbALFA fluorophores, while the signal of all other epitope tags was amplified using polyclonal secondary antibodies. Without further optimization, NbALFA yielded a remarkably linear signal over at least three orders of magnitude (Fig.2d) and was able to detect as little as 100pg of target protein. The detection limit was thus ~10-times better than observed for all other epitope tags.

**Fig. 2.**
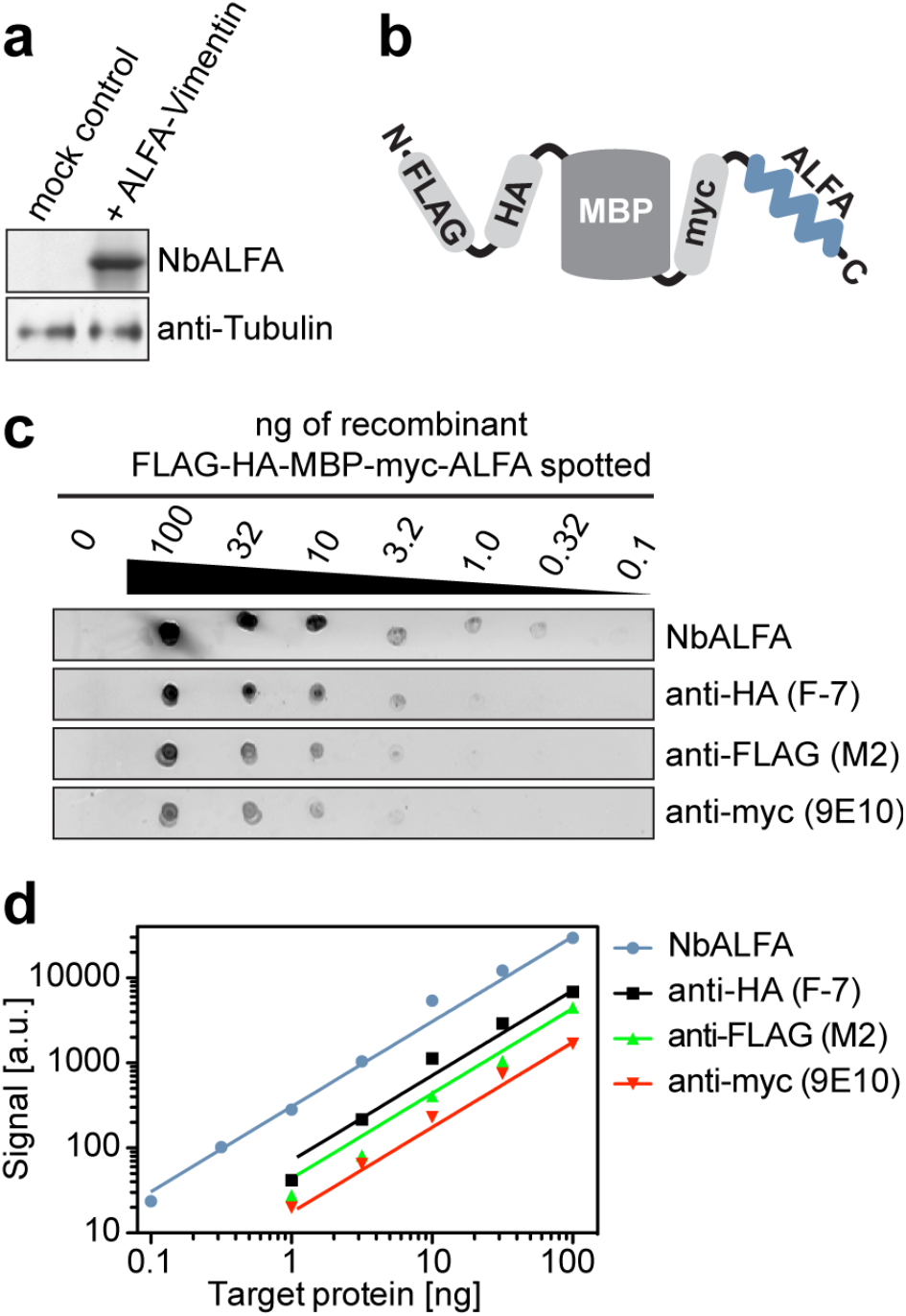
Detection of ALFA-tagged target proteins by fluorescent Western blot. **a;** Lysates from COS-7 cells transfected with ALFA-Vimentin or a control plasmid were analyzed by SDS-PAGE and Western blot. The blot was developed with NbALFA directly coupled to IRDye800CW or a mouse anti-Tubulin followed by FluoTag-X2 anti-Mouse IRDye680RD. Complete lanes are shown in Fig.S4a. **b;** Sketch of the *E. coli* maltose-binding protein (MBP) multi-tag fusion used for experiments shown in panels **c** and **d**. **c;** Dilution series of the protein sketched in **b** were spotted onto nitrocellulose membranes. Established monoclonal antibodies (M2, 9E10 and F-7) were used together with an anti-mouse IgG coupled IRDye800CW to detect the FLAG^®^, myc and HA-tags respectively. The ALFA-tag was detected using NbALFA coupled to IRDye800CW. Fig.S4b shows the complete experiment including internal controls. **d;** Double-logarithmic plot showing quantification of signals obtained in **c**. Lines represent linear fits to the obtained values. Even without signal amplification by a secondary antibody, signals obtained using NbALFA were 3-to >10-times stronger than by established reagents recognizing the other epitope tags. At the same time, detection with NbALFA was 10-fold more sensitive and showed an excellent linearity over ~3 orders of magnitude.

### Structure of the NbALFA bound to the ALFA peptide

To better understand the interaction between NbALFA and the ALFA tag, we solved the crystal structure of NbALFA in complex with the ALFA peptide (Fig.3, Table S2). NbALFA adopts a canonical IgG fold^30^ comprising two β-sheets connected by a disulfide bond. The paratope accepting the ALFA peptide extends from the nanobody’s N-terminal cap to the side formed by the five-stranded β-sheet. It is mainly built from complementarity determining regions (CDRs^15^) 2 and 3 and, interestingly, also involves residues within the five-stranded β-sheet forming a hydrophobic cavity. As a result, the ALFA peptide is oriented parallel to the central axis of NbALFA. The ALFA peptide forms an α-helical cylinder with a length of ~2nm and a diameter of ~1.3nm, that is stabilized by a complex network of intramolecular interactions (Table S3). In addition, the peptide forms multiple polar and hydrophobic contacts with NbALFA (Fig.3b-e, Table S4).

**Fig. 3:**
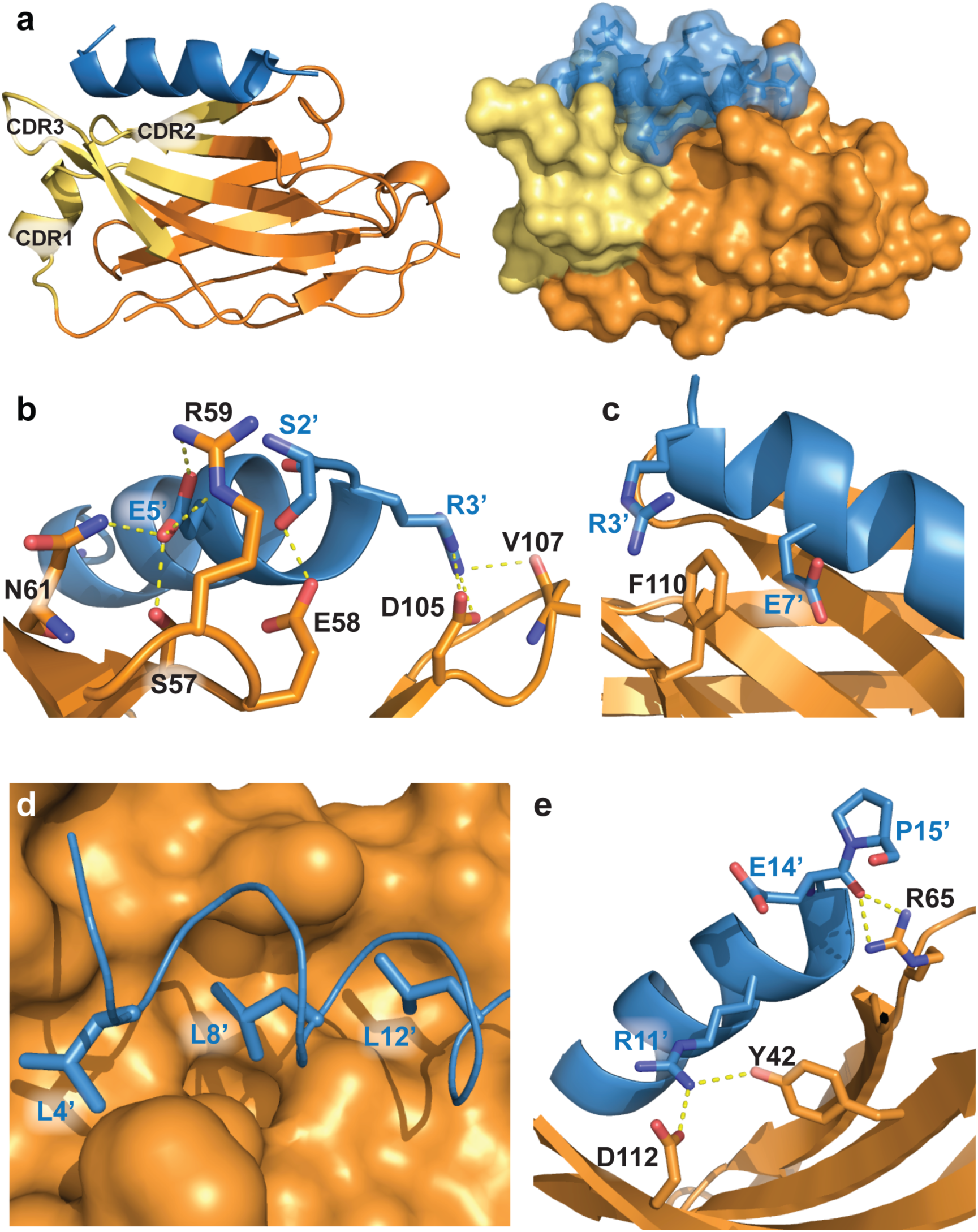
Structure of the NbALFA-ALFA peptide complex. **a;** View on the NbALFA-ALFA peptide complex illustrated as cartoon (left) or surface representation (right). NbALFA: orange with CDRs 1-3 colored in yellow. ALFA peptide in blue. For both molecules, the N-terminal is oriented left, the C-terminal right. **b;** Polar interactions within the N-terminal region of the ALFA peptide. ALFA peptide residues are denoted by an apostrophe. S2’ and E5’ form hydrogen bonds with CDR2 (S57, E58, R59 and N61). R3^’^ reaches out to CDR3 and interacts with the backbone of V107 and the side chain of D105. **c;** R3’ and E7’ sandwich F110 on the nanobody, forming a cation-Pi interaction (reviewed in^29^); Illustration of a hydrophobic cluster (L4’, L8’ and L12’) facing the nanobody’s hydrophobic cavity. **e;** Polar interaction near the C-terminal of the ALFA peptide. The backbone of E14’ forms hydrogen bonds with R65 while the side chain of R11’ interacts with D112 and Y42 on the five-stranded β-sheet. Interestingly, Y42 has been described as a conserved residue in nanobodies of this particular architecture^8^.

### Capture of ALFA-tagged target proteins using ALFA Selector resins

For biochemical purifications, we site-specifically immobilized NbALFA on an agarose-based resin (Fig.4a). The resulting resin efficiently and strongly bound an ALFA-tagged GFP variant (shGFP2^31^): Even after 1h competition with an excess of free ALFA peptide at 25°C, most of target protein remained on the resin (Fig.4b, red solid line). In line with these observations, SPR assays indicated that the affinity of NbALFA to an ALFA-tagged target was ~26 pM (Fig.S5). We therefore called the NbALFA-charged resin “ALFA Selector^ST^” (for ***S***uper-***T***ight). To allow for an efficient competitive elution of ALFA-tagged target proteins, we intended to weaken NbALFA. For that, we followed a rational approach based on the NbALFA-ALFA peptide structure: NbALFA variants featuring single or combined amino acid exchanges in spatial proximity to the ALFA peptide were individually tested for their binding and elution properties. The successfully weakened binder NbALFA^PE^ (for **P**eptide **E**lution) carries several mutations with respect to NbALFA, thereby removing specific interactions of NbALFA with the ALFA peptide. As a consequence, the affinity of NbALFA^PE^ to a fusion protein harboring a C-terminal ALFA-tag was reduced to ~11 nM (Fig.S5). As intended, a NbALFA^PE^-charged resin (ALFA Selector^PE^) efficiently and stably captured shGFP2-ALFA while allowing an efficient release within ~15-20 minutes by competition with free ALFA peptide (Fig.4b,c). Similar elution kinetics were found when the ALFA-tag was placed between two folded domains, while elution of ALFA-shGFP2 from ALFA Selector^PE^ was slightly quicker (Fig.S6). Remarkably, in the absence of competing peptide, spontaneous elution of all target proteins from both ALFA Selector variants was insignificant (Fig.4b, c and Fig.S6).

**Fig. 4:**
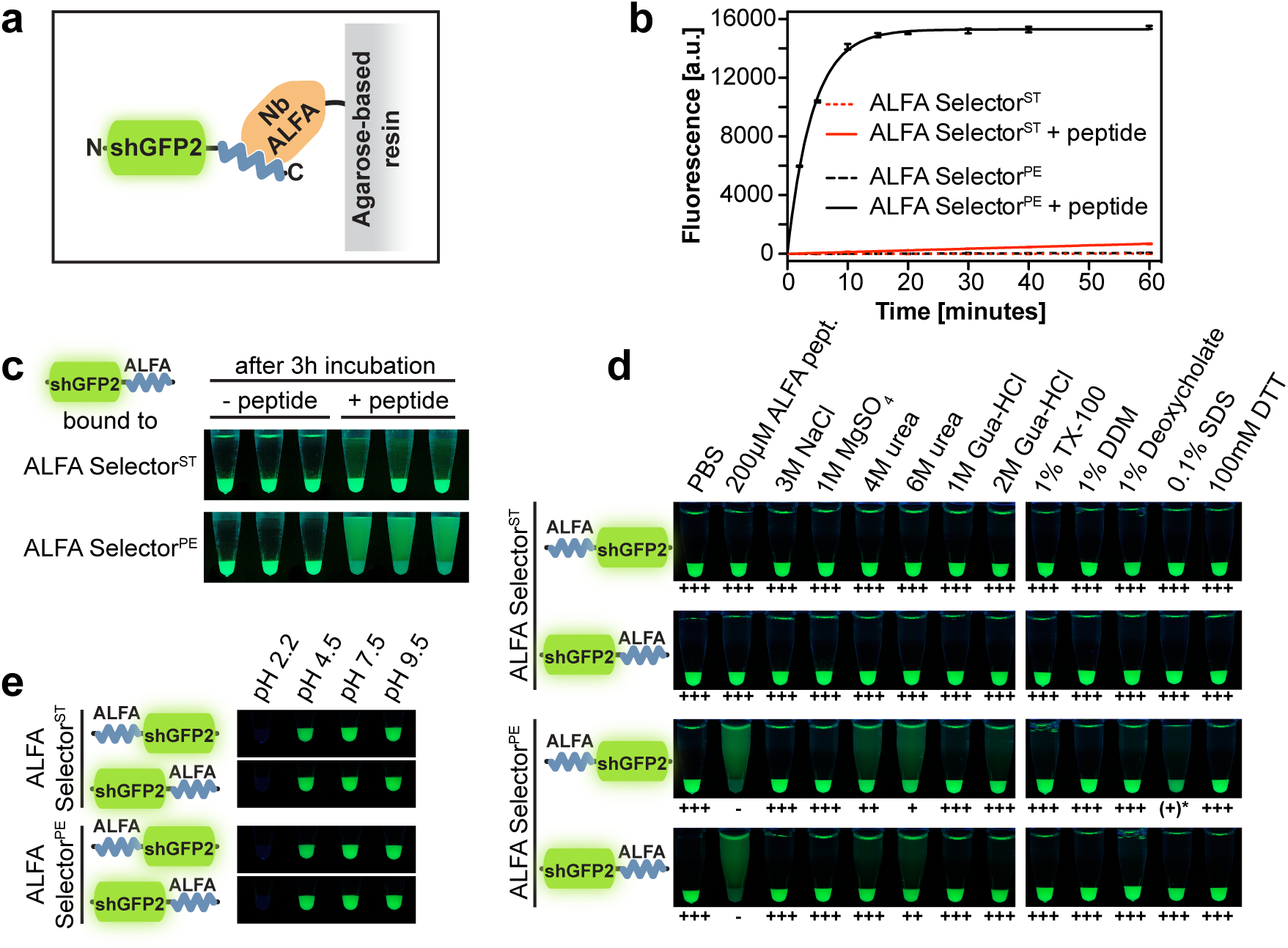
Immunoprecipitation of ALFA-tagged proteins using ALFA Selector resins. **a;** Sketch of shGFP2-ALFA bound to a resin coupled to a NbALFA. **b and c;** ALFA Selector^ST^ or ALFA Selector^PE^ were charged with shGFP2 harboring a C-terminal ALFA-tag. To estimate off-rates, the resins were incubated with an excess of free ALFA peptide at 25°C. Control reactions were carried out without peptide. shGFP2 released from the resin was quantified using its fluorescence. The graph (**b**) shows mean fluorescence readings as well as standard deviations (n=3). Lines represent fits to a single exponential. A photo was taken upon UV illumination after 3h of elution (**c**). **d;** Resistance towards stringent washing steps. Both ALFA Selector variants were charged with either ALFA-shGFP2 or shGFP2-ALFA and incubated with a 10-fold volume of the indicated substances for 1h at 25°C with shaking. Without further washing steps, photos were taken upon UV illumination after sedimentation of the beads. **e;** Resistance towards various pH: Similar to **d**, here, however, the resins were washed to remove non-bound material after incubating at indicated pH for 30min. Photos were taken after re-equilibration in PBS to allow for recovery of the GFP fluorescence.

### Stringent washing and pH resistance

We next performed stringent washing steps on both ALFA Selectors variants bound to either ALFA-shGFP2 or shGFP2-ALFA (Fig.4d). At 25°C, all substrate–resin interactions were completely resistant to significant concentration of salt, Guanidinium-HCl, non-denaturing detergents or reducing agents. A small fraction of substrate was released from ALFA Selector^PE^, but not from ALFA Selector^ST^, upon treatment with up to 6M urea.

In a similar assay, the loaded ALFA Selector resins were exposed to buffers adjusted to various pH (Fig.4e). The interaction was stable at pH7.5 or 9.5, and only slightly affected at pH4.5. However, even after neutralization, both ALFA Selector resins remained completely non-fluorescent when washed with 100mM Glycin at pH2.2. The eluted material, in contrast, successfully recovered its fluorescence at neutral pH (not shown), indicating that acidic elution with Glycin at pH2.2 is possible even from ALFA Selector^ST^.

### Pull-down of ALFA-tagged target proteins from complex lysates

To address the specificity of the ALFA Selector resins in native pull-downs, we spiked *E. coli* or HeLa lysates prepared in PBS with recombinant ALFA-shGFP2 (Fig.5a, left lane). The target protein specifically bound to both ALFA Selector variants but not to a control resin without coupled nanobody. As expected, ALFA-shGFP2 efficiently eluted from ALFA Selector^PE^ using 200µM of ALFA peptide in PBS. In contrast, significant elution from ALFA Selector^ST^ was only observed upon treatment with SDS sample buffer. Importantly, pull-downs from both, *E. coli* (Fig.5a) and HeLa lysates (Fig.5b) were highly specific: After peptide elution from ALFA Selector^PE^ essentially all visible bands could be attributed to the input protein, and even in the SDS eluate, the number and strength of detectable impurities originating from lysate proteins were very low.

**Fig. 5.**
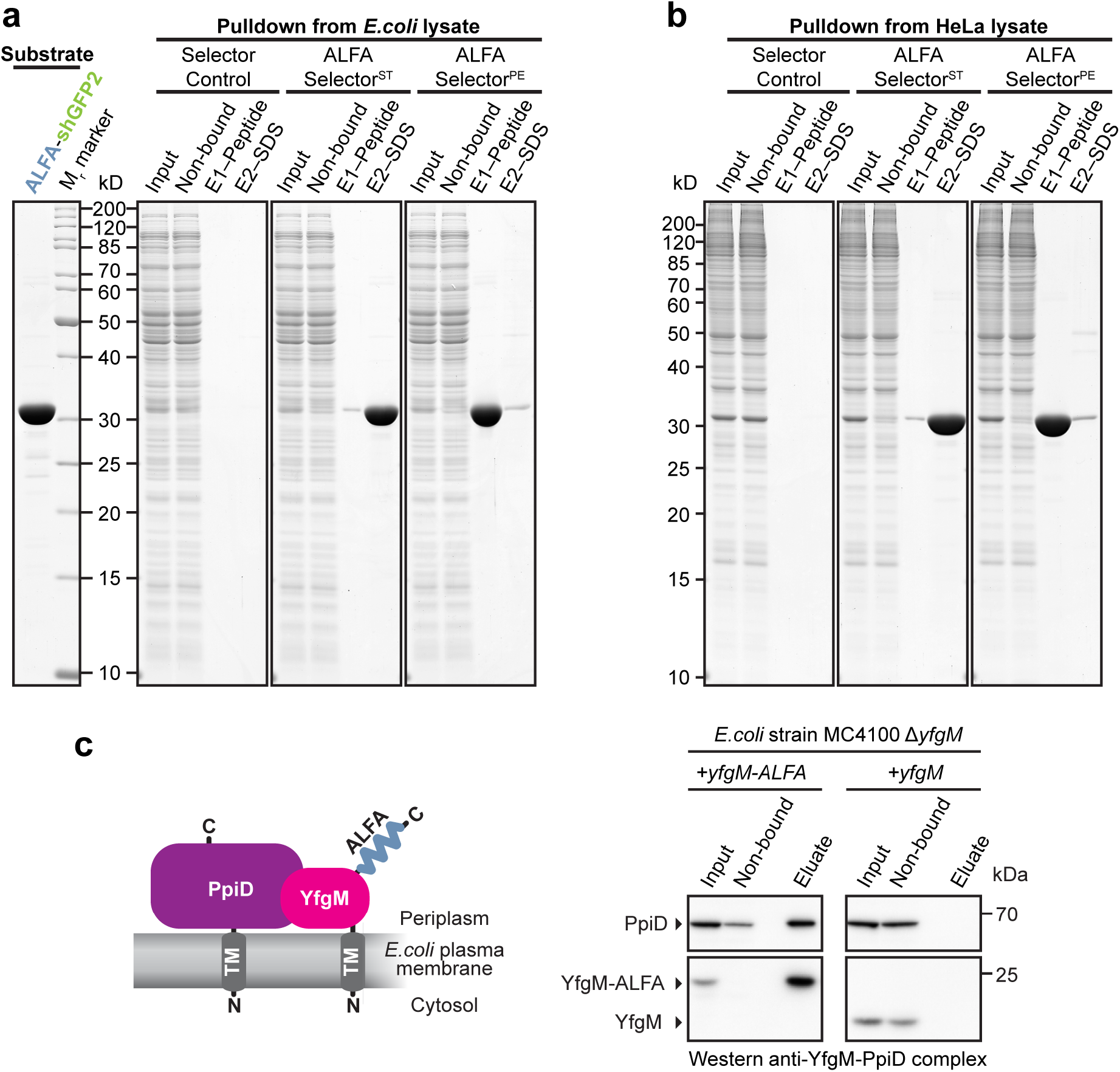
One-step affinity purifications using ALFA Selector resins. **a and b;** *E. coli* (**a**) or HeLa (**b**) lysates blended with purified ALFA-tagged shGFP2 (**a**, left lane) were incubated with ALFA Selector^ST^, ALFA Selector^PE^ or an analogous resin without immobilized sdAb (Selector Control). After washing with PBS, the resins were incubated with 200µM ALFA peptide for 20min before eluting remaining proteins with SDS sample buffer. Indicated fractions were analyzed by SDS-PAGE and Coomassie staining. Eluate fractions correspond to the material eluted from 1µL of resin. **c;** Native pull-down of an *E. coli* inner membrane protein complex. Left: Sketch of the target protein complex. Right: Detergent-treated lysates from a Δ*yfgM* strain complemented with either C-terminally ALFA-tagged (left panel) or untagged YfgM (right) were incubated with ALFA Selector^PE^. After washing with PBS, bound proteins were eluted using 200µM ALFA peptide. Samples corresponding to 1/800 of the input and non-bound material or 1/80 of eluate fractions were resolved by SDS page and analyzed by Western blot. ALFA Selector^PE^ specifically immunoprecipitated the native protein complex comprising ALFA-tagged YfgM and its interaction partner PpiD. In the control reaction (no ALFA tag on YfgM), both proteins were absent in the eluate. Complete blots in Fig.S7.

In a more delicate co-immunoprecipitation experiment, we tried to pull-down the *E. coli* YfgM-PpiD inner membrane protein complex^32^ under native conditions (Fig.5c). For this, either wild-type YfgM or YfgM-ALFA was expressed in a Δ*yfgM* strain under the control of its endogenous promoter^32^. ALFA Selector^PE^ was able to specific pull-down the YfgM-PpiD complex in a detergent-resistant manner from the lysate containing YfgM-ALFA. The native membrane protein complex could be recovered from ALFA Selector^PE^ by peptide elution under physiological conditions showing that the ALFA-tag together with ALFA Selector^PE^ resin is perfectly suited for native pull-downs of challenging (membrane) protein complexes.

### Isolation of live lymphocytes

An envisioned application for the ALFA Selector^PE^ is the specific enrichment of cells under physiological conditions. This may be particularly interesting e.g. for the generation of chimeric antigen receptor-modified T (CAR-T) cells, the precursors of which are usually obtained from blood^33^. To investigate if the ALFA system can be applied to enrich live blood cells, human peripheral blood mononuclear cells (PBMCs) were passed through an ALFA Selector^PE^ column pre-charged with an ALFA-tagged nanobody recognizing CD62L, a surface marker typically present on naïve T cells^34^ (Fig.6a). After washing, bound cells were eluted using ALFA peptide, stained with antibodies recognizing CD62L, the pan T cell marker CD3 and the pan B cell marker CD19, and analyzed by FACS (Fig.6). Total PBMCs served as a control. Using this strategy, CD62L^+^ lymphocytes were enriched from 71.8 to 97.7% (Fig.6b). In addition, we confirmed that the vast majority of ALFA peptide-eluted cells were CD3-positive T cells, while B cells represented a minor population of the isolated cells (Fig.6c).

**Figure 6.**
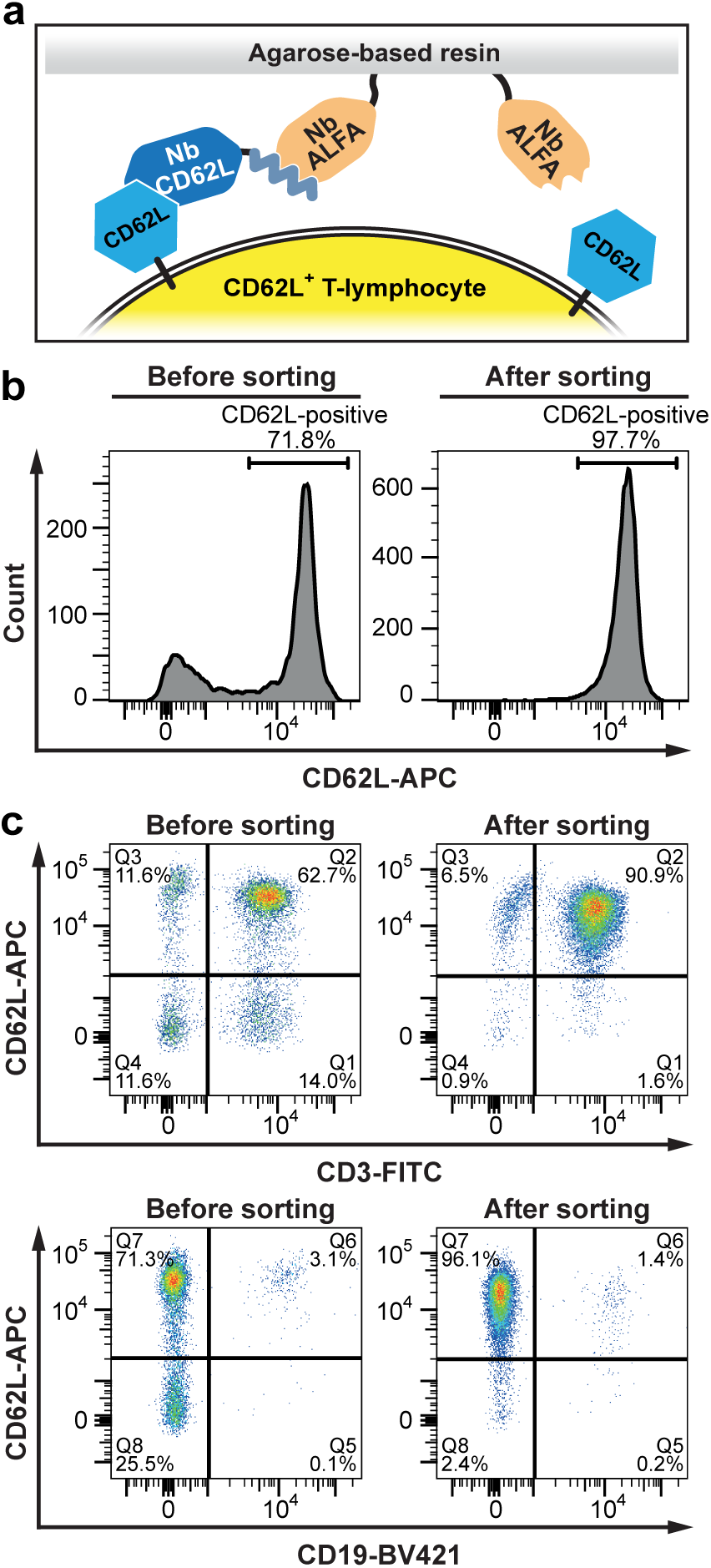
Isolation of naïve lymphocytes using an ALFA-tagged nanobody recognizing CD62L. Total human PBMCs were left untreated (Before sorting) or isolated using an ALFA Selector^PE^ resin loaded with an ALFA-tagged anti-human CD62L nanobody (After sorting). A sketch of the affinity purification strategy is shown in (**a**). Cells were stained with an anti-CD62L antibody and analyzed by flow cytometry (**b**). The same cells as in (**b**) were stained with antibodies directed against CD3, CD19 and CD62L, and analyzed by flow cytometry (**c**). A forward scatter/side scatter gate was set on lymphocytes in all analyses.

## Discussion

In the current study, we report the characterization of the ALFA system comprising the ALFA-tag, a novel and highly versatile epitope tag, and two dedicated nanobodies. The mainly rational approach allowed us to equip the ALFA system with favorable features for a broad spectrum of applications. When selecting the ALFA-tag sequence, we not only made sure that the tag is small, monovalent, devoid of lysines, hydrophilic without carrying any net charge and absent within the proteome of relevant model organisms, but also that it would adopt a stable α-helical structure in solution. As a result, the ALFA-tag is by design highly specific (Fig.5), insensitive to amine-reactive fixatives (Fig.1c) and well tolerated by the tagged target proteins (Figs.1, S1-S3). The ALFA-tag could thus even be used for purification of a labile membrane protein complex (Fig.5c). However, as for any other tag, specific effects on given target proteins have to be analyzed on a protein-to-protein basis.

The ALFA tag with its comprehensive feature set contrasts to existing tags, which mostly fail to fulfill one or more parameters (Table S1). For instance, large structured tags like GFP can affect the tagged protein localization and function^4,18^. On the other hand, intrinsically disordered small tags can adopt multiple conformations with unpredictable effects. The HA-tag, e.g., can be cleaved by caspases in mammalian cells^35^. In the worst case, tags may even target the POI for protein degradation^36^. Aldehyde-containing fixatives used for light and electron microscopy modify lysine-containing epitope tags like the myc-tag, which may impair antibody-based detection. Other tags (e.g. the FLAG^®^-tag) carry significant net charges and may thus affect the POI’s physiological electrostatic interactions. Tags like myc-tag^9^, the HA-tag^10^, the Spot-tag^®11,12^, C-tag^14^ and the Inntags^37^ are derived from proteins present in model organisms. Therefore, the respective binders can also recognize the endogenous host proteins by default. In contrast to the C-tag^14^ that only works at a POI’s C-terminal, the ALFA-tag works at all accessible locations within the target protein without interfering with the protein’s localization or topology (Figs.S1-S3).

For binding the ALFA-tag we developed nanobodies^15^, because they are small, monovalent and robust, can easily be modified by genetic means and recombinantly produced in various organisms. They can therefore be site-specifically immobilized or quantitatively modified with fluorophores^24^ or oligonucleotides. In comparison to conventional antibodies, nanobodies are thus superior for conventional and advanced microscopy^11,17^ (Fig.1e,f) since they can find more target epitopes in crowded areas^22^, localize the fluorophore closer to the target protein and avoid artificial clustering of the POI^16,21^. Interestingly, NbALFA folds even within eukaryotic cells (Fig.1g) and can thus be used as an intrabody^7^ for detecting or manipulating ALFA-tagged proteins *in vivo*^7,8,26^.

Nanobodies often fail to detect proteins in Western blot applications as they typically recognize three-dimensional epitopes. NbALFA, however, enables highly sensitive Western blots, suggesting that the ALFA-tag’s α-helical structure refolds efficiently after SDS removal. Despite its monovalent binding mode and without further signal amplification, NbALFA significantly outperformed established anti-epitope tag tools regarding absolute signal intensity and detection limit (Fig.2). We expect a superior performance also in other high sensitivity applications like ELISA or microarray assays. Due to its resistance to amine-reactive fixatives (Fig.1d), we even envision applying the ALFA system for immuno-electron microscopy and nanoSIMS^38^.

The binding of NbALFA to the ALFA-tag is exceptionally strong, which could be explained after solving the crystal structure of the NbALFA-ALFA peptide complex (Fig.3). First, the peptide binds to NbALFA in a α-helical conformation and has an unusually high propensity to form a stable α-helix also in solution^23^. This may be explained by multiple intramolecular side-chain interactions within the peptide (Table S3). As a result, no energy needs to be spent in forming the required structure during complex formation, which would otherwise disfavor complex formation. Second, the peptide forms multiple specific contacts with the nanobody (Table S4) along the whole length of the 13 amino acid ALFA core sequence (SRLEEELRRRLTE). In sum, nearly all residues of this sequence are involved in binding to NbALFA and/or stabilizing the α-helical peptide conformation. Due to the compact structure of the complex, the maximal displacement between an ALFA-tagged target and a fluorophore attached to NbALFA is well below 5nm and can be reduced to <3nm by choosing adequate positions for fluorescent labeling. In order to minimize the potential influences of neighboring sequences on the ALFA-tag conformation, we placed the ALFA core sequence between prolines acting as “insulators”. Using this approach, the interaction of NbALFA with the ALFA-tag is largely independent of its localization within the tagged protein and is efficiently recognized when placed at either terminus of the POI or even in between two separately-folded domains.

The strong interaction of NbALFA and the ALFA-tag is ideal for imaging applications, highly sensitive detection of target proteins, and purification of extremely low-abundant proteins from dilute lysates or under conditions where harsh washings with chaotropic agents are required. The slow dissociation, however, precludes a competitive elution of ALFA-tagged proteins under physiological conditions within a reasonable time frame. Based on the crystal structure of the NbALFA-ALFA peptide complex (Fig.3), we site-specifically mutated the nanobody to increase its off-rate. A resin displaying the mutant nanobody (ALFA Selector^PE^) allows for native purifications of proteins and protein complexes from various lysates under physiological conditions by peptide elution (Figs.4, 5). ALFA Selector^PE^ could even be applied for the selective enrichment of CD62L-positive lymphocytes from PBMC preparations (Fig.6). We believe that this technique can easily be transferred to the highly validated recombinant Fab and scFv fragments that are currently used for cell isolation approaches and similar purposes^39^, or to novel nanobodies recognizing surface markers that can easily be equipped with an ALFA tag. Our new technology can therefore contribute to current advances in biomedical research and therapy including the CAR-T technology^33^.

The novel ALFA tag system stands out by its exceptionally broad applicability. Using the ALFA system, a single transgenic cell line or organism harboring an ALFA-tagged target protein is sufficient for a wealth of different applications including (super-resolution) imaging, *in-vivo* manipulation of proteins, *in-vitro* detection by Western blot or even native pull-down applications aiming at detecting specific interaction partners or at isolating specific cell populations. The wide range of applications of the ALFA system provides the scientific community with a novel and highly versatile tool, which will facilitate future scientific breakthroughs.

## Acknowledgements

We thank Alexandra Lück and Verena Pape for excellent technical assistance, the scientists at BM14.2, HZB, Berlin for their support during X-ray diffraction data collection, Jonas Ries for SMAP software support and Florian Schueder, Alexander Auer, and Maximilian T. Strauss for support with DNA-PAINT experiments. We further thank Mihalea Serpe for directing our attention to potential *in-vivo* applications, Blanche Schwappach for her generous support regarding affinity determinations, and Eugenio F. Fornasiero and Silvio Rizzoli for comments on the manuscript. FO and SSI were supported by the DFG through Cluster of Excellence Nanoscale Microscopy and Molecular Physiology of the Brain (CNMPB). This work was partially supported by grants from the Swedish Research Council (2014-5667) and the Swedish Cancer Society to P.S.. T.S. acknowledges support from the DFG through the Graduate School of Quantitative Biosciences Munich (QBM). R.J. is supported by the DFG through the Emmy Noether Program (DFG JU 2957/1-1), the SFB 1032 (Project A11) and by the ERC Starting Grant (MolMap; 680241), by the Max Planck Society and the Center for Nanoscience (CeNS).

## Author contributions

SF and FO conceived the project. SF, HG and FO were involved in experimental design, nanobody selection and development of the ALFA system. SF, HG, MK and FO performed and analyzed most of the experiments. AR initially characterized the NbALFA^PE^ mutant. MMC and PS solved and analyzed the NbALFA-ALFA peptide structure. NE helped with the cell isolations and performed the FACS analysis. SSI contributed with critical reagents for DNA-PAINT and affinity measurements. TS and RJ performed and analyzed DNA-PAINT experiments. SF, HG, MK and FO wrote the manuscript. All authors read and commented on the manuscript.

## Materials & Correspondence

For plasmid requests please write to

## Data Availability

The atomic coordinates and structure factors (code 6I2G) have been deposited in the Protein Data Bank (http://www.pdb.org). Primary data of graphs shown in Figures 4b, S2, S5 and S6 are available in the Source Data file.

## Online Methods

**Table M1:** Plasmids used in this study.

| Transfection |  |  |
| --- | --- | --- |
| Identifier | Encoded protein | Origin/Citation |
| pNT1112 | ALFA-FLAG<sup>®</sup>-Vimentin | This study |
| pNT1067 | Tom70-EGFP-ALFA | This study |
| pNT1066 | pDisplay-HA-EGFP-ALFA-myc-TM | This study |
| pNT1004 | NbALFA-mScarlet-I | This study |
| pEGFP-N1 | EGFP | Clontech |
| pNT0942 | EGFP-ALFA | This study |
| pNT0941 | ALFA-EGFP | This study |
| Bacterial expression |  |  |
| Identifier | Encoded protein | Origin/Citation |
| pNT1208 | His<sub>14</sub>-bdSUMO-FLAG<sup>®</sup>-HA-MBP-myc-ALFA | This study |
| pNT1111 | His<sub>14</sub>-bdSUMO-NbALFA | This study |
| pNT1177 | ALFA-shGFP2-His<sub>6</sub> | This study |
| pNT1050 | His<sub>14</sub>-bdSUMO-shGFP2-ALFA | This study |
| pNT1063 | His<sub>14</sub>-bdSUMO-ALFA-shsfGFP | This study |
| pNT1129 | pSC YfgM-ALFA | This study |
| pSC-yfgM | *yfgM* (native promoter in pUA66) | Götzke *et. al.*<sup>32</sup> |
| pNT1196 | His<sub>14</sub>-bdSUMO-TST-bdNEDDD8-ALFA | This study |
| pNT1298 | His<sub>14</sub>-bdSUMO-NbCD62L-ALFA | This study |

### E.coli strains

*E. coli* MC4100 Δ*yfgM* Δ*ppiD*^32^

**Table M2:** Antibodies.

| Antibody | Supplier | Order No |
| --- | --- | --- |
| FluoTag<sup>®</sup>-X2 anti-ALFA<br>AbberiorSTAR635P<br>(NbALFA-Ab635P) | NanoTag Biotechnologies | N1502-Ab635P |
| FluoTag<sup>®</sup>-X2 anti-ALFA IRDye800CW | NanoTag Biotechnologies | N1502 (Custom) |
| FluoTag<sup>®</sup>-X2 anti-Mouse IRDye680RD | NanoTag Biotechnologies | N1202 (Custom) |
| anti-FLAG<sup>®</sup> M2 | Sigma | F1804 |
| anti-myc 9E10 | SynapticSystems | #343 011 |
| anti-HA F-7 | SantaCruz | #sc-7392 |
| Goat anti-rabbit IRDye680RD | Li-COR | #925-68071 |
| Goat anti-mouse IRDye800CW | Li-COR | #925-32210 |
| anti-Tubulin | SynapticSystems | #302 211 |
| polyclonal serum recognizing MBP | SynapticSystems |  |
| goat anti-rabbit IgG HRP-conjugate | Dianova | #GAR/IgG(H+L)/PO |
| anti-YfgM/PpiD | *Götzke et. al.*<sup>32</sup> |  |
| anti-human CD3, clone UCHT1, FITC | BioLegend | 300405 |
| anti-human CD19, clone HIB19, BV421 | BioLegend | 302233 |
| anti-human CD62L, clone DREG-56,<br>APC | BD Biosciences | 559772 |

### Protein expression and purification

All recombinant proteins were expressed under the control of the Tac-promoter from expression vectors with ColE1 origin that confer resistance to Kanamycin.

The MBP fusion protein harboring multiple epitope tags, ALFA-shGFP2, TwinStrepTag-bdNEDDD8-ALFA, NbCD62L-ALFA and non-tagged NbALFA were expressed as N-terminal His_14_-bdSUMO fusions. For protein expression, *E. coli* was cultured in Terrific broth (TB) supplemented with 0.3mM IPTG for 14-16 hours at 23°C. After harvest, *E. coli* cells were lysed in LS buffer (50mM Tris/HCl pH7.5, 300mM NaCl) supplemented with 15mM imidazole/HCl pH7.5 and 10mM DTT, and purified by binding to Ni(II)-chelate beads. After extensive washing, proteins were eluted by on-column-cleavage with bdSENP1 as described before40,41. ALFA-shGFP2-His_6_ and His_14_-bdSUMO-ALFA-shsfGFP were expressed and purified in a similar fashion; Elution was, however, performed using 250mM Imidazole in buffer LS. For affinity determinations and binding studies from complex lysates, substrate proteins were additionally purified via size exclusion chromatography on a Superdex200 10/30 column (GE Healthcare). NbALFA harboring N- and C-terminal ectopic cysteines was expressed and purified as described for non-tagged NbALFA above. Labeling with maleimide-activated fluorophores was performed as previously described elsewhere^24^.

### Selection of specific sdAb clones by affinity purification of B lymphocytes

1mL of T-Catch resin (IBA Lifesciences) was washed with B cell isolation buffer (PBS pH7.4, 1% BSA, 1mM EDTA) and incubated with saturating amounts of a TwinStrepTag-bdNEDDD8-ALFA fusion protein for 30min rolling at RT. The resins were cleared from excess bait protein by extensively washing with B cell isolation buffer. 100mL of blood sample was taken from alpaca immunized with ALFA peptide and immediately incubated with 5000 IU/mL heparin (Sigma) to prevent clotting. From the fresh blood (less than 4h past sampling) PBMCs were isolated using Ficoll-Paque PLUS (GE Healthcare). To remove residual serum, PBMCs were washed three times consecutively with B cell isolation buffer. PBMCs were passed over the loaded T-Catch resin three times before washing the resins with 10 CV B cell isolation buffer. Bound B cells were eluted from the resins by incubating 2µM NEDP1^40,41^ for 30min at RT. From the eluted B cells an sdAb-specific cDNA library was amplified by a multistep nested RT-PCR and cloned into a bacterial expression vector. 96 single clones were tested by ELISA for expression of ALFA-reactive sdAbs.

### Crystallization and structure determination

For crystal formation, non-tagged NbALFA was incubated with a 1.2-fold excess of ALFA peptide (Acetyl-PSRLEEELRRRLTEP-Amide; Genscript) for 60min on ice. The complex was concentrated to ~25mg/mL and separated from the excess peptide by gel filtration on a Superdex75 10/30 column (GE Healthcare) equilibrated with 20mM HEPES pH7.4, 100mM NaCl, 10% glycerol.

Crystals of the NbALFA-ALFA peptide complex were found in the F2 condition of the JCSG+ screen (Molecular Dimensions). This hit was further optimized using the Hampton Research Additive Screen. A crystal suitable for data collection was found in a drop containing 100nL of precipitant solution (100mM sodium citrate pH6.0, 2M ammonium sulfate, 4% v/v tert-Butanol) and 100nL of protein solution (10.1mg/ml NbALFA-ALFA peptide complex, 20mM HEPES pH7.4, 100mM NaCl, 10% glycerol) in a sitting-drop crystallization plate. X-ray diffraction data collection was performed at BL14.2 at the BESSY II electron storage ring operated by the Helmholtz-Zentrum Berlin, Germany^42^. The data were processed using DIALS^43^, molecular replacement was performed with Phaser^44^ using the structure with PDB code 5VNV as an atomic model. The structure was built using Phenix Autobuild^45^ and Coot^46^. The structure was refined using Refmac5^47^.

### Affinity determinations

All surface plasmon resonance (SPR) binding experiments were performed on a Reichert SR 7500DC biosensor instrument at 20°C and a flow rate of 40µL/min on HC1000m SPR sensor chips (Xantec Biotechnologies) in PBS buffer containing 0.002% (v/v) Tween-20, pH 7.4. Data were analyzed using TraceDrawer v1.7.1. The HC1000m sensorchip was activated with 1-ethyl-3-3(3-dimethylaminiopropyl)-carbodiimide and N-hydroxysuccinimide (EDC-NHS) according the manufacturers instructions. The ligand was pre-diluted to 5µM in concentration buffer (50mM Sodium acetate pH 5.0) and it was injected over the activated chip surface at a flow rate of 15µl/min to a final concentration of 150µRIU (TST-NEDD8-ALFA) and 650µRIU (GFP-ALFA) response units (RIU: refractive index units; response recorded). The chip surface was subsequently inactivated with 1M ethanolamine pH 8.5. The chips with immobilized ligands were equilibrated with PBS pH 7.4 containing 0.002% (v/v) Tween-20 (PBS-T). The analytes (NbALFA^PE^, or NbALFA) were serially diluted in PBS-T and injected over the chip surface. Association was followed for 4.5min and dissociation was followed for 15min for NbALFA^PE^ and accordingly 45min for NbALFA. For each analyte, two buffer injections were performed as a reference (buffer reference). The recorded binding was double-referenced against an empty reference channel recorded in parallel and the buffer reference. The surface was regenerated after each analyte injection by injecting 100mM Glycine pH 2.2, 150mM NaCl for 2.5min. To accurately determine the dissociation constant of NbALFA, the decrease in response was followed for >7h.

### Transfection of 3T3, COS-7 or HeLa cells

For immunofluorescence experiments, 3T3, COS-7 or Hela (Leibniz Institute DSMZ) cells were transiently transfected with 0.2 to 1µg of plasmids listed in Table M1, using the PolyJet transfection kit (SignaGen) or lipofectamine 3000 (Thermo Fisher Scientific, cat. no. L3000-001) according to the manufacturer’s recommendations. 3T3 and COS-7 cells were seeded on 12-well plates and HeLa in 8-well chambered coverglass (ibidi, cat. no. 80827). Cells were incubated with transfection reagents for 24-48 hours. For co-expression experiments, plasmid DNA was premixed in a 1:1 ratio and further processed as described above.

### Microscopy Setups

Epifluorescence images were obtained with an Axio Observer Z1, (Carl Zeiss GmbH) equipped with a 20x dry lens or a 40x oil immersion lens. Confocal and STED microscopy were acquired using an inverse 4-channel Expert Line easy3D STED setup (Abberior Instruments GmbH, Göttingen, Germany). The setup was based on an Olympus IX83 microscope body equipped with a plan apochromat 100× 1.4 NA oil-immersion objective (Olympus). 640nm excitation laser (Abberior Instruments GmbH) pulsed at 40 MHz was used. Stimulated depletion was achieved with a 775nm STED laser (Abberior Instruments GmbH) pulsed at 40 MHz with an output power of ~1.250 W. Fluorescence signal was detected using APD detectors (Abberior Instruments GmbH). The operation of the setup and the recording of images were performed with the Imspector software, version 0.14 (Abberior Instruments GmbH). DNA-PAINT imaging was carried out on an inverted Nikon Eclipse Ti microscope (Nikon Instruments) with the Perfect Focus System, applying an objective-type TIRF configuration with an oil-immersion objective (Apo SR TIRF 100×, NA 1.49, Oil). Two lasers were used for excitation: 561nm (200mW, Coherent Sapphire) or 488nm (200mW, Toptica iBeam smart). The laser beam was passed through a cleanup filter (ZET488/10x or ZET561/10x, Chroma Technology) and coupled into the microscope objective using a beam splitter (ZT488rdc or ZT561rdc, Chroma Technology). Fluorescence light was spectrally filtered with two emission filters (ET525/50m and ET500lp for 488nm excitation and ET600/50 and ET575lp for 561nm excitation, Chroma Technology) and imaged on a sCMOS camera (Andor Zyla 4.2) without further magnification, resulting in an effective pixel size of 130nm after 2×2 binning. Camera Readout Sensitivity was set to 16-bit, Readout Bandwidth to 540 MHz.

### Epifluorescence and STED microscopy imaging

Transiently transfected 3T3 or COS-7 cells were fixed in either 4% paraformaldehyde (PFA) (*w*/*v*) or 2% glutaraldehyde (GA) (*v*/*v*) for 30min at room temperature (RT). Alternatively, fixation was performed in ice cold methanol for 15min at −20°C. Cells were blocked and permeabilized in PBS containing 10% normal goat serum (*v*/*v*) and 0.1% Triton-X 100 (*v*/*v*) for 15min at RT. Fluorescently labeled NbALFA (FluoTag^®^-X2 anti-ALFA AbberiorStar635P, NanoTag Biotechnologies #N1502-Ab635P-L) was diluted 1:500 in PBS containing 3% normal goat serum and 0.1% Triton-X 100 (*v*/*v*). The cells were incubated in this staining solution for 1h at RT and subsequently washed 3 times for 5min with PBS. To stain the nucleus, DAPI (0.4µg/mL) was included in one of the PBS washing steps. Coverslips were mounted on cover-slides using Mowiol solution, dried at 37°C and imaged. Constructs expressed at the cell-surface were co-stained with anti-FLAG^®^ M2 (primary antibody, Sigma, F1804) and FluoTag^®^-X2 anti-mouse IgG Atto488 (secondary nanobody, NanoTag Biotechnologies #N1202-At488-L) diluted 1:1000 and 1:500 respectively, in PBS containing 3% normal goat serum and 0.1% Triton-X 100 (*v*/*v*).

### DNA-labeling of NbALFA, DNA-PAINT imaging and data analysis

NbALFA was first coupled to a single stranded DNA as described before^48^. In brief, NbALFA bearing ectopic N- and C-terminal cysteines was reduced on ice with 5mM of tris(2-carboxyethyl)phosphine (TCEP, Sigma-Aldrich, #C4706) for 2h. After removing excess TCEP using 0.5ml Amicon Ultra spin filters with a molecular weight cut-off (MWCO) of 10 kDa (Merck, #UFC500324), the nanobody was conjugated with a 50-fold molar excess of maleimide-DBCO crosslinker (Sigma-Aldrich, #760668) at 4°C overnight. Using a 10 kDa MWCO Amicon spin filter, the excess of crosslinker was removed and the nanobody was further incubated with 10 fold molar excess of a single stranded DNA containing an azide group on its 5′-end and an Atto488 fluorophore on its 3′ end (P3 docking strand: 5’-Azide-TTTCTTCATTA-Atto488-3’ obtained from Biomers.net GmbH) for 2h at room temperature. Excess of DNA was removed by using a size exclusion chromatography column (Superdex^®^ Increase 75, GE Healthcare) on an Äkta pure 25 system (GE Healthcare).

For staining, cells were prefixed with 0.4% (*v/v*) Glutaraldehyde and 0.25% (*v/v*) Triton X-100 in PBS at pH7.2 for 90s. The main fixation was performed with 3% glutaraldehyde in PBS for 15min. Afterwards the sample was reduced with 1 mg/mL sodium borohydride (Carl Roth, #4051.1) for 7min and washed 4x (1x fast, 3× 5min) with PBS. Blocking and additional permeabilization was performed for 90min in 3% (*w/v*) BSA + 0.2% (*v/v*) Triton X-100 in PBS. Next, NbALFA conjugated to the P3 docking strand was added to the sample to a final concentration of 5µg/mL in dilution buffer (3% *w/v* BSA in PBS) and incubated for 1h at RT. The sample was washed 3x for 5min in PBS and then incubated for 10min with 1:5 diluted fiducial markers, 90-nm gold particles (Cytodiagnostics, #G-90-100), residual gold was washed away with PBS. Cells were kept at 4°C until they were used for imaging.

For DNA-PAINT imaging, transfected cells were screened for a certain phenotype with 488nm laser excitation at 0.01kW/cm^2^. After acquisition of the 488nm channel, the excitation was switched to 561nm, focal plane and TIRF angle were readjusted and imaging was subsequently performed using ~2.5kW/cm^2^ 561nm laser excitation. P3-imager strand (5’-GTAATGAAGA-Cy3b-3’) concentration was chosen to minimize double-binding events. ALFA-tag imaging was performed using an imager concentration of 1nM P3-Cy3b in Buffer C (PBS + 500mM NaCl). All imaging was performed in 1×PCA (Sigma-Aldrich, #37580)/1×PCD (Sigma-Aldrich, #P8279)/1×Trolox (Sigma-Aldrich, #238813) in Buffer C and cells were imaged for 40,000 frames at 100ms camera exposure time. 3D imaging was performed using a cylindrical lens in the detection path as previously reported^49^. Raw data movies were reconstructed with the Picasso software suite^25^. Drift correction was performed with a redundant cross-correlation and gold particles as fiducials. Final images obtained had a localization precision of <10nm calculated via a nearest neighbor analysis^50^. Rendering was performed via the recently published SMAP software.

### Impact of ALFA-tags on the localization of EGFP

Transiently transfected 3T3 cells were imaged using an epifluorescence microscope (Axio, Zeiss) equipped with a 40× 1.3 oil lens. For cells transfected with either pCMV ALFA-EGFP, pCMV EGFP-ALFA, or pEGFP-N1, 107-133 cells were imaged in a total of six to seven individual images. For each individual image, cells were grouped and counted according to the localization of EGFP (“slightly nuclear”, “equally distributed”, “other”). The fraction of cells in each group was statistically analyzed using Student’s t-test.

### Western Blots with COS-7 lysates

Transfected cells from a confluent 10 cm petri dish were washed with PBS and lysed in 2mL SDS sample buffer. Lysates were resolved by SDS-PAGE and transferred to a nitrocellulose membrane. After blocking with 5% milk powder in TBS-T, membranes were incubated with mouse anti-tubulin (Synaptic Systems #302 211; 1:1000 dilution) followed by a FluoTag^®^-X2 anti-Mouse IgG IRDye680 (NanoTag Biotechnologies #N1202; 1:1000 dilution) and FluoTag-X2 anti-ALFA IRDye800 (NanoTag Biotechnologies #1502; 1:1000 dilution). Membranes were scanned using Odyssey CLx (Li-COR).

### Dotblot assay

A serial dilution of MBP fused to FLAG^®^-, HA-, myc- and ALFA-tags was prepared in PBS pH7.4, 0.1µg/mL BSA. 1µL of each dilution was spotted on nitrocellulose membranes. Membranes were blocked and washed with 5% milk powder in TBS-T. Established monoclonal antibodies (anti-FLAG^®^ M2, Sigma #F1804; anti-myc 9E10 SynapticSystems #343 011; anti-HA F-7, SantaCruz #sc-7392) were used in combination with a secondary goat anti-mouse IgG IRDye800CW (Li-COR #925-32210, dilution 1:500) to detect FLAG^®^, myc and HA-tag, respectively. The ALFA-tag was detected using a FluoTag^®^-X2 anti-ALFA (NanoTag Biotechnologies #N1502) directly coupled to IRDye800CW. All primary antibodies and the nanobody were used at 2.7nM final concentration. Detection of MBP by a rabbit polyclonal serum recognizing MBP (SynapticSystems) and an anti-rabbit IgG IRDye680RD (Li-COR #925-68071) served as an internal loading control. Membranes were scanned using Odyssey CLx (Li-COR). Quantifications were performed using ImageStudioLight (Li-COR).

### Off-rate assays

20µL ALFA Selector^ST^ or ALFA Selector^PE^ (NanoTag Biotechnologies) was saturated with the respective recombinant target protein. After washing 4x with PBS, the beads were suspended in a 10-fold excess of PBS containing 200µM free ALFA peptide and mixed at 25°C. Control reactions were carried out without peptide. At indicated time points, specific elution from the beads was quantified using the GFP fluorescence released into the supernatant (Q-Bit 3.0; Thermo-Fischer Scientific). Three independent experiments were performed in parallel. Mean values, standard deviations and exponential fits were calculated using GraphPad Prism 5.0. Photographic pictures were taken upon UV illumination using a Nikon D700 camera equipped with a 105mm macro lens (Nikon).

### Resistance towards stringent washing and pH

15µL of ALFA Selector^ST^ or ALFA Selector^PE^ saturated with either ALFA-shGFP2 or shGFP2-ALFA were washed with PBS and incubated with 100µL of the indicated substances for 60min at RT. Photos were taken after sedimentation of the beads upon UV illumination. To assay for pH resistance, the same beads were incubated with 150mM NaCl buffered to various pH (100mM Glycin-HCl, pH2.2; 100mM Na-Acetate pH4.5, 100mM Tris-HCl pH7.5, 100mM Carbonate pH9.5) for 30min at RT. The resin was washed twice with the same buffer. Photos were taken after equilibrating several times with PBS.

### One-step affinity purifications from *E. coli* and HeLa lysates

To obtain defined input materials for pull-down experiments from *E. coli* or HeLa lysates, respective mock lysates were blended with 3µM of purified ALFA-tagged shGFP2. 1mL of each lysate/substrate mixture was incubated with 25µL of ALFA Selector^ST^ or ALFA Selector^PE^ for 1h at 4°C. An analogous resin without immobilized sdAb (Selector Control) served as a specificity control. After washing 3 times with 600µL of PBS, the resins were transferred into MiniSpin columns (NanoTag Biotechnologies). Excess buffer was removed by centrifugation (3000x g, 30s) before incubating twice for 10min at RT with 50µL each of 200µM ALFA peptide in PBS. Proteins remaining on the beads were afterwards eluted with SDS sample buffer. 0.5µL (*E. coli*) or 1.5µL (HeLa) of input and non-bound fractions were resolved by SDS-PAGE (12%) and Coomassie staining. Shown eluate fractions correspond to the material eluted from 1µL of the respective resins.

### Native pull-down of the *E. coli* YfgM-PpiD inner membrane protein complex

A *yfgM* deletion strain was complemented with either C-terminally ALFA-tagged or untagged YfgM expressed from a pSC-based low-copy vector under control of the endogenous promoter. Membrane protein complexes were solubilized from total lysates prepared in buffer LS (50mM Tris pH7.5, 300mM NaCl, 5mM MgCl_2_) using 1% DDM for 1h on ice^51^. Both lysates were incubated with 20µL of ALFA Selector^PE^ resin for 1h at 4°C on a roller drum. After washing in PBS containing 0.3% DDM, bound proteins were eluted under native conditions by sequentially incubating twice with 50µL PBS containing 200µM ALFA peptide. Samples corresponding to 1/800 of the input and non-bound material or 1/80 of eluate fractions were resolved by SDS-PAGE. Analysis was performed by Western blotting using a polyclonal rabbit antiserum raised against the YfgM-PpiD complex^32^ followed by an HRP-conjugated goat anti-rabbit IgG (Dianova). Blots were developed using the Western Lightning Plus-ECL Kit (Perkin Elmer) and imaged using a LAS 4000 mini luminescence imager (Fuji Film).

### Preparation of human PBMCs

Human peripheral blood mononuclear cells (PBMCs) were obtained from fresh blood using standard density gradient centrifugation. Briefly, 60mL of fresh blood was diluted with 40mL of phosphate-buffered saline (PBS) supplemented with 1mM EDTA and placed on top of a layer of CELLPURE Roti-Sep 1077 (Carl Roth) in 50mL LEUCOSEP tubes (Greiner Bio-One) and centrifuged at 800 × g for 20min at room temperature. Subsequently, the PBMC-containing layer was collected and washed five times in cold PBS + EDTA to remove platelets.

### Isolation of CD62L-positive lymphocytes

Approximately 2 × 10^7^ PBMCs were passed by gravity flow through an ALFA Selector^PE^ resin loaded with an ALFA-tagged anti-human CD62L nanobody, followed by extensive washing with PBS supplemented with 1mM EDTA and 1% (w/v) bovine serum albumin. Subsequently, bound cells were eluted in the same buffer containing 200µM ALFA peptide.

### Flow cytometry

For flow cytometric analysis, roughly 1 × 10^6^ cells per sample were stained with monoclonal antibodies to human CD3 (clone UCHT1) coupled to FITC (BioLegend), CD19 (clone HIB19) coupled to BV421 (BioLegend) and CD62L (clone DREG-56) coupled to APC (BD Biosciences). After washing with PBS supplemented with EDTA and BSA, cells were measured using an LSRII cytometer (BD Biosciences). Data were analyzed using FlowJo (v10) software (FlowJo, LLC, Ashland, OR, USA).

