## Supplementary material for "A rationally designed and highly versatile epitope tag for nanobody-based purification, detection and manipulation of proteins": Fig.S1



Nanobody, sdAbs, epitope tag, affinity, immunoprecipitation, native elution, super-resolution microscopy, STED, DNA-PAINT, CAR-T, lymphocytes, cell isolation

### Supplementary Figures

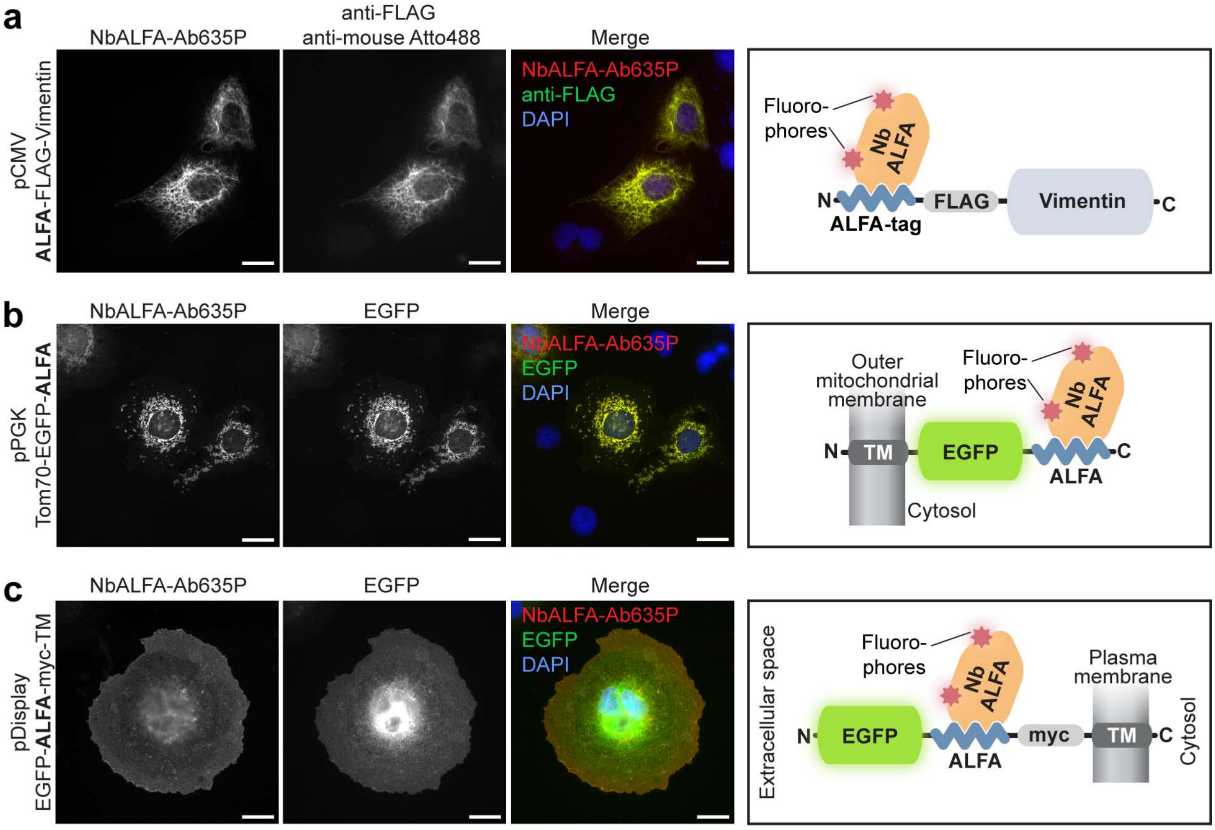


**Figure S1 – related to Figure 1: ALFA-tagged proteins can be detected by fluorescently labeled NbALFA regardless of the localization of the tag within the fusion protein.**

COS-7 cells were transfected with constructs encoding proteins fused to an ALFA-tag at their N-terminus (ALFA-FLAG®-Vimentin; **a)** C-terminus (Tom70-EGFP-ALFA; **b**), or within individual protein-domains (EGFP-ALFA-myc-TM; **c**). Cells were fixed with 4% PFA and stained as indicated. For experiments shown in panels **a** and **b**, cells were permeabilized with 0.1% TritonX-100; for panel **c**, cells were stained under non-permeabilizing conditions. TM: transmembrane domain. Scale bars: 20µm. Sketches illustrate the topology of substrates and detection by fluorescently labeled NbALFA (NbALFA-Ab635P).

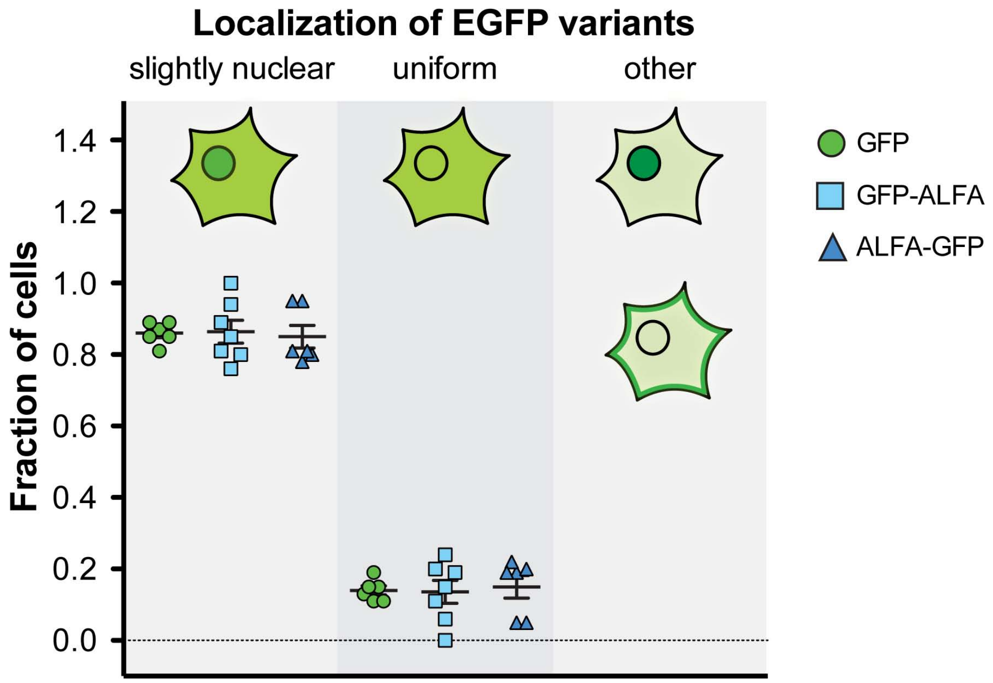


**Figure S2 – related to Figure 1: N- or C-terminal ALFA-tag fusions show the correct intracellular localization.**

3T3 cells were transiently transfected with EGFP fusions harboring N- or C-terminal ALFA tags. Non-tagged EGFP from pEGFP-N1 served as a control. The localization of the respective EGFP variants was analyzed on 6-7 individual images for each construct. Between 120 and 130 cells were imaged per construct and the localization of EGFP was analyzed. In general, each EGFP construct displayed a distribution across the cytosol and the nucleus. Cells were distributed into three groups (”slightly nuclear”, “uniform” and “other”) according to the observed nucleocytoplasmic localization of EGFP. Standard deviations were calculated from values obtained from individual images. Differences between the localizations of tagged and non-tagged EGFP variants were statistically not significant (Student’s t-test).

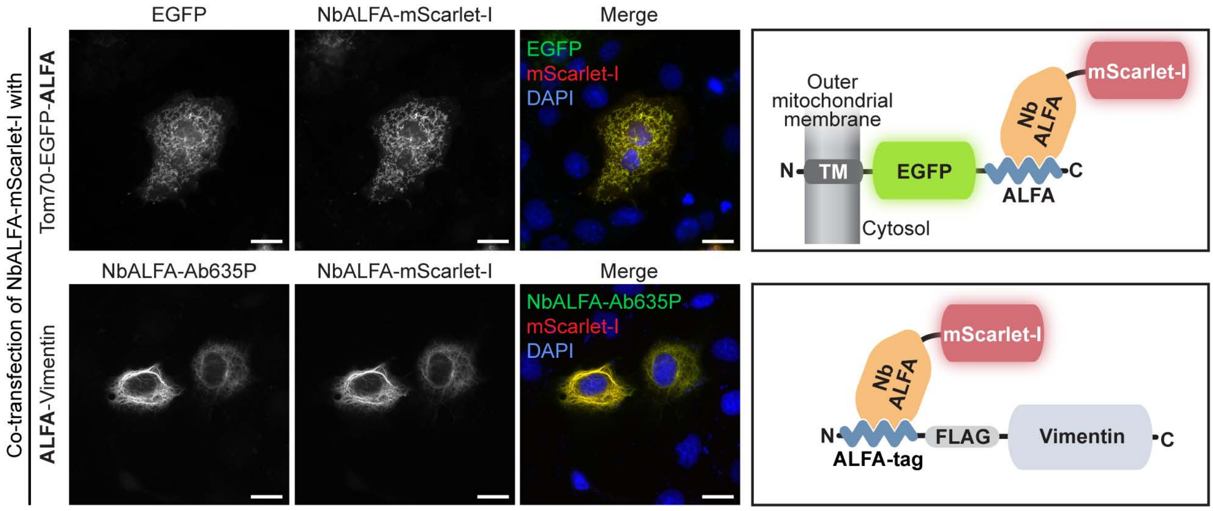


**Figure S3 – related to Figure 1: Target protein detection using NbALFA expressed the cytoplasm of mammalian cells.**

COS-7 cells were co-transfected with an NbALFA-mScarlet-I fusion and indicated ALFA-tagged target proteins. Target proteins were detected either using the intrinsic EGFP fluorescence (TOM70-EGFP-ALFA) or by immunofluorescence using NbALFA-Ab635P (ALFA-Vimentin). In parallel, NbALFA-mScarlet-I was detected by its red fluorescence. The excellent co-localization shows that NbALFA expressed in the cytoplasm of mammalian cells can be used for targeting ALFA-tagged proteins in living cells. Scale bars: 20µm.

The lower panel recapitulates data shown in Figure 1g and is repeated here for a better comparison.

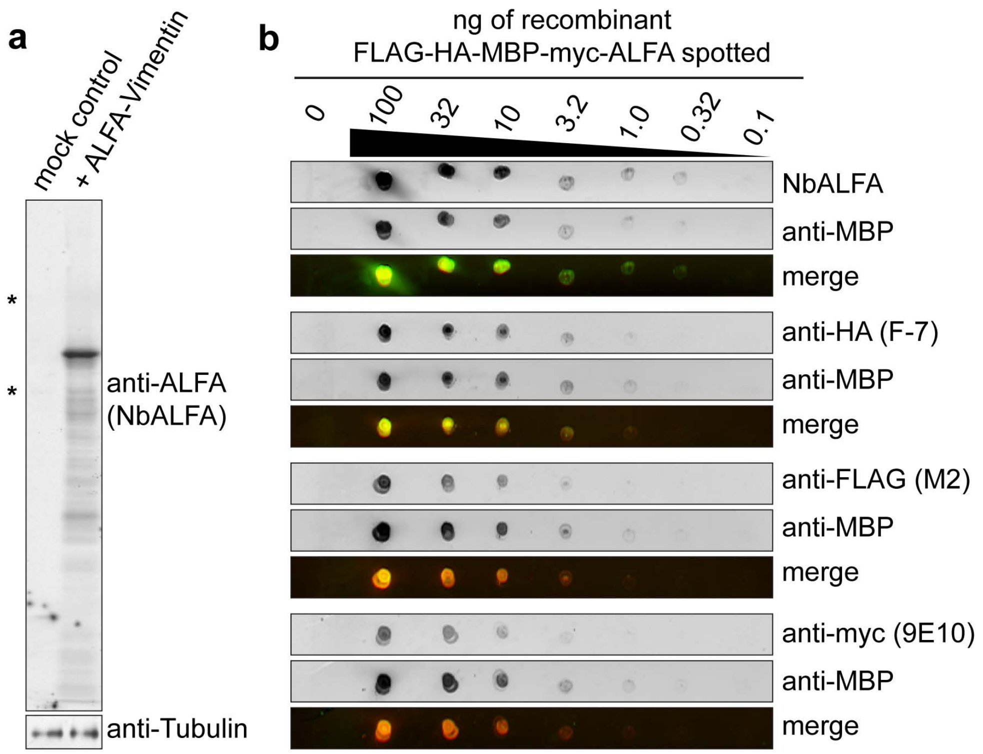


**Figure S4 –related to Figure 2: Highly sensitive detection of ALFA-tagged target proteins by Western blot using fluorescently labeled NbALFA.**

**a**, Complete lanes of the Western blot shown in Figure 2a. Note that in the absence of any vector encoding an ALFA-tagged protein, only very minor bands (*) can be detected using NbALFA directly labeled with IRDye800CW**. b**, In addition to the data presented in Figure 2c, detection of MBP by a combination of rabbit polyclonal serum recognizing MBP and an anti-rabbit IgG IRDye680RD (Li-Cor #925-68071) is shown as an internal loading control. Individual signals are in gray levels and, colored overlays show MBP signals in red and signals corresponding to epitope tags in green.

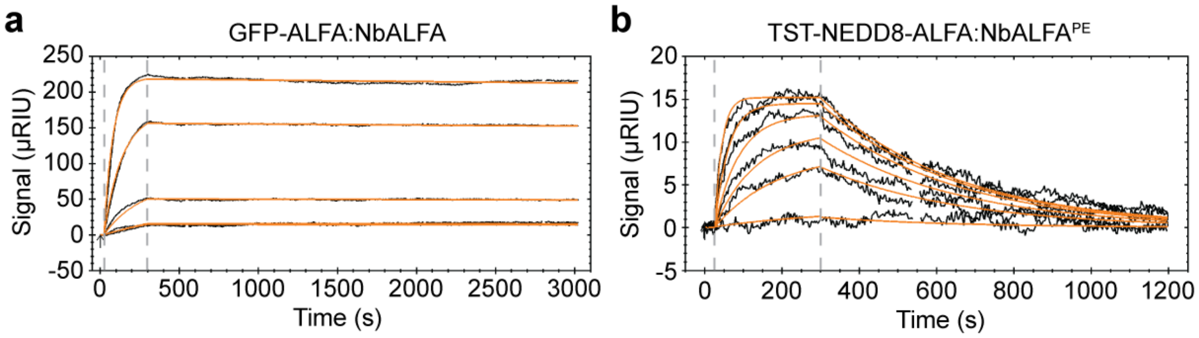


c

| Nanobody | Target | $k_{on}$ ($M^{-1}s^{-1}$) | $k_{off}$ ($s^{-1}$) | $K_D$ (nM) |
|---|---|---|---|---|
| **NbALFA** | GFP-ALFA | 3.6 (± 0.1) x $10^{5}$ | 9.4 (± 0.2) x $10^{-6}$ | 0.026 (± 0.001) |
| **NbALFA$^{PE}$** | TST-NEDD8-ALFA | 2.7 (± 0.4) x $10^{5}$ | 2.9 (± 0.5) x $10^{-3}$ | 11.0 (± 0.6) |

**Figure S5: Affinity of NbALFA and NbALFA$^{PE}$ for ALFA-tagged target proteins.**

**a,** Representative SPR sensogram for NbALFA$^{ST}$ binding to an ALFA-tagged target protein. A serial dilution of analyte (NbALFA$^{ST}$; 50nM, 25nM, 12.5nM, 6.25nM, 3.13nM) was injected over a sensorchip surface coated with GFP-ALFA. Black lines: data recorded; orange lines: fits used to analyze data. **b,** Representative SPR sensogram for NbALFA$^{PE}$ binding to an ALFA-tagged target protein. A serial dilution of analyte (NbALFA$^{PE}$; 200nM, 100nM, 50nM, 25nM, 12.5nM, 6.25nM) was injected over a sensorchip surface coated with TST-NEDD8-ALFA. Black lines: data recorded; orange lines: fits. **c,** Summary of affinity data obtained from 3 (NbALFA) or 4 (NbALFA$^{PE}$) independent experiments, respectively. Standard deviations are given in brackets.

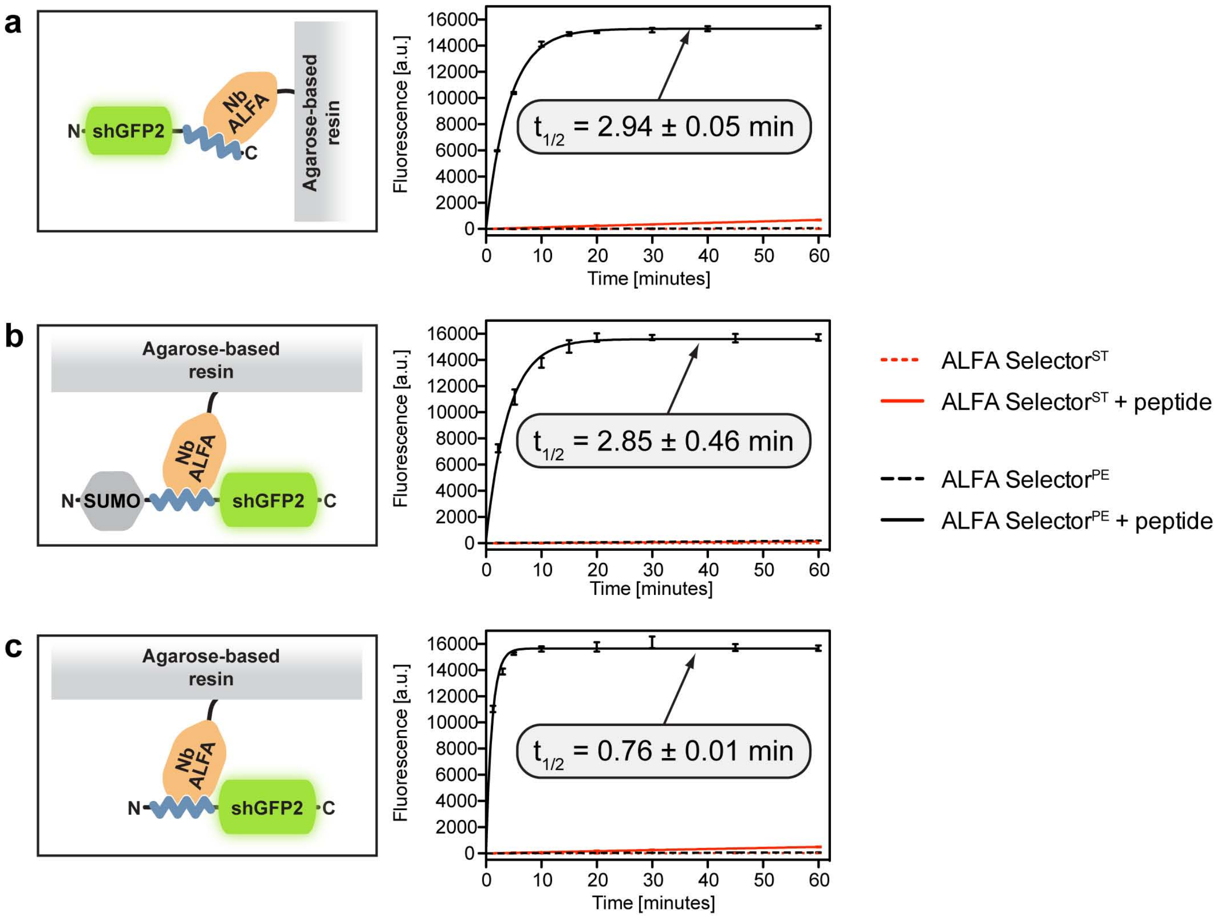


**Figure S6 – related to Figure 4b: Peptide elution of ALFA-tagged GFPs from ALFA Selector resins.**

20µL ALFA Selector$^{ST}$ (presenting NbALFA) or ALFA Selector$^{PE}$ (presenting NbALFA$^{PE}$) were charged with shGFP2-ALFA (**a**), bdSUMO-ALFA-shGFP2 (**b**) or ALFA-shGFP2 (**c**) as examples of C-terminal, in between folded domains and N-terminal ALFA-tagged POI respectively. After washing with PBS, the beads were suspended in a 10-fold excess of PBS containing 200µM free ALFA peptide and gently mixed at 25°C. Control reactions were carried out without peptide. At indicated time points, specific elution from the beads was quantified using the GFP fluorescence released into the supernatant. Shown are mean fluorescence readings of three experiments as well as standard deviations for each time point. Lines represent fits to a single exponential. Half times are given for peptide elution from ALFA Selector$^{PE}$ only. For all substrate proteins, peptide elution from ALFA Selector$^{ST}$ was inefficient even after prolonged incubation. In the absence of ALFA peptide, the ALFA-tagged target proteins remained tightly bound to both resins. Note that peptide elution of shGFP2 harboring an N-terminal ALFA tag from ALFA Selector$^{PE}$ is quicker as compared to substrates harboring the ALFA tag at internal or C-terminal positions. Panel **a** recapitulates data shown in Figure 4b and is repeated here to allow a direct comparison.

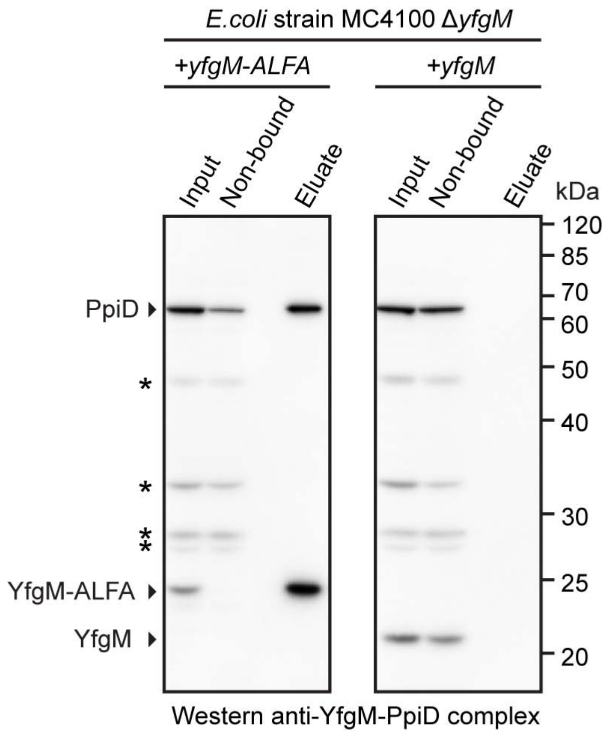


**Figure S7 – related to Figure 5c: Pull-down of a native *E. coli* YfgM-PpiD inner membrane protein complex using the ALFA Selector (complete blots).**

A *yfgM* deletion strain was complemented with either C-terminally ALFA-tagged (left panel) or untagged YfgM (right panel; control reaction) expressed from a low-copy vector. Membrane protein complexes were solubilized from total lysate using DDM. Complexes containing YfgM-ALFA were purified in a single step using ALFA Selector$^{PE}$ affinity resin and eluted under native conditions using 200 µM ALFA peptide. Samples corresponding to 1/800 of the input and non-bound material or 1/80 of eluate fractions were resolved by SDS-PAGE and analyzed by Western blot. A rabbit serum raised against the YfgM-PpiD complex recognized both, PpiD and YfgM, in the input fractions. ALFA Selector$^{PE}$ specifically immunoprecipitated the native protein complex comprising ALFA-tagged YfgM and its interaction partner PpiD. In the control reaction (no ALFA tag on YfgM), both proteins were absent in the eluate. The figure recapitulates data shown in Figure 5c, here, however, complete membranes are shown.

### Table S1: Properties of common epitope tag systems

| | Parameter | FLAG-tag[1] | HA-tag[2] | myc-tag[3] | Twin-Strep-tag[4] | polyHis-tag[5,6] | EPEA/C-tag[7] | Spot-tag[8,9] | ALFA tag |
|---|---|---|---|---|---|---|---|---|---|
| Properties of tag | Sequence | DYKDDDDK | YPYDVPDYA | EQKLI-SEEDL | WSHPQFEK-(GGGS)$_2$-GGSA-WSHPQFEK | His$_{3-10}$ | EPEA | PDRVRA-VSHWSS | (P)SRLEEELR RRLTE(P) |
| | Size (amino acids) | 8 | 9 | 10 | 28 | 3-10 | 4 | 12 | 14-15 |
| | Mass (kD) | 1.1 | 1.1 | 1.2 | 2.9 | 0.4-1.4 | 0.5 | 1.4 | ~1.8 |
| | Charge at pH7.0 | -3 | -2 | -3 | 0.2 | 0.3-1 | -2 | 1.1 | 0 |
| | pKi | 3.5 | 0 | 3.5 | 8.4 | 14 | 0 | 12.1 | 8.1 |
| | Physical size (nm) | 2.2 | 2.5 | 2.8 | >6 | 0.8-2.8 | 1.2 | 3.3 | 2.0 |
| | Water solubility | + | poor | + | + | poor | + | + | + |
| | Structured in solution | – | – | – | – | – | - | – | +[10] |
| | Stable | (+)[11] | (+)[12] | + | + | + | n.d. | + | + |
| | Resistant to aldehyde modification[(a)] | – | + | – | – | + | + | + | + |
| | Unique within model organisms | + | – | – | (+) [(b)] | (–) [(c)] | – | (–) [(d)] | + |
| Properties of binder | Name | M1, M2, M5 | F-7, 12CA5 | 9E10 | StrepTactin-XT | $Ni^{2+}$/$Co^{2+}$-chelate | NbSyn2 | Spot-nanobody | NbALFA |
| | Type | mAb | mAb | mAb | Protein | Inorganic | sdAb | sdAb | sdAb |
| | Size (kD) | ~150 | ~150 | ~150 | ~60 | n.a. | ~15 | ~15/30 [(e)] | ~15 |
| | No. of polypeptides | 4 | 4 | 4 | 4 | n.a. | 1 | 1 | 1 |
| | No of binding sites | 2 | 2 | 2 | 2 | 1 | 1 | 1-2 [(e)] | 1 |
| | Affinity | n.d. | n.d. | n.d. | ~60 pM [(f)] | n.d.[(g)] | >190 nM | ~6 nM | 26 pM |
| | Genetically accessible | – | – | – | + | – | + | + | + |
| Appli-cations | Protein purification | + | + | + | + | + | + | + | + |
| | Imaging | + [(h)] | + [(h)] | + [(h)] | n.d. | (–) [(i)] | – | (+) [(e)] | + |
| | *In-vivo* applications | – | – | – | – | – | – | – | + |

Abbreviations: n.a.: not applicable; n.d.: no data available; mAb: monoclonal antibody; sdAb: single-domain antibody

(a) Fixation by amine-reactive fixatives and cross-linkers; deduced from sequence
(b) Binder also recognizes biotinylated proteins
(c) Binder recognizes endogenous proteins with multiple accessible histidines.
(d) Binder also recognizes endogenous β-catenin
(e) Binder needs to be dimerized for high-profile imaging applications[8]

(f) https://www.iba-lifesciences.com/tl_files/ProteinProductionAssays/5-Immobilization/DynamicBiosensors-Application-Note-StrepTactinXT-switchSENSE.pdf
(g) Depends on chelate and polyHis-tag used
(h) Multiple tags used in tandem for optimal performance[13–16]
(i) Rarely used for imaging in combination with Tris-NTA derivatives[17]

**Table S2. Data collection and refinement statistics.**

| **Data collection** | |
|---|---|
| Space group | C 1 2 1 |
| Cell dimensions | |
| *a*, *b*, *c* (Å) | 101.24, 32.38, 64.73 |
| a, b, g (°) | 90.00, 147.22, 90.00 |
| Resolution (Å) | 1.5 |
| $R_{sym}$ or $R_{merge}$ | 0.063 (0.225) |
| *I* / s*I* | 12.70 (4.26) |
| Completeness (%) | 95.2 (94.0) |
| Redundancy | 5.6 (5.4) |
| **Refinement** | |
| Resolution (Å) | 50.33 - 1.5 |
| No. reflections | 17569 / 865 |
| $R_{work}$ / $R_{free}$ | 0.165 / 0.200 |
| No. atoms | |
| Protein | 1067 |
| Ligand/ion | 28 |
| Water | 118 |
| *B*-factors | |
| Protein | 11.31 |
| Ligand/ion | 25.68 |
| Water | 18.56 |
| R.m.s. deviations | |
| Bond lengths (Å) | 0.0129 |
| Bond angles (°) | 1.857 |

**Table S3.** List of intramolecular side-chain interactions within the ALFA peptide.

| **Residue [a]** | **Interaction** | **Distance (Å)** |
|---|---|---|
| E7'-R10' | Salt bridge | 2.67 |
| E7'-R11' | Salt bridge | 2.77 |
| R9'-T13' | H-bond | 3.07 |
| L4'-L8'-L12' | Hydrophobic array | |

[a] ALFA peptide residues are marked by an apostrophe.

**Table S4.** List of intermolecular interactions between NbALFA and the ALFA peptide.

| **ALFA peptide residue [a]** | **NbALFA residue** | **Interaction** | **Distance (Å)** |
|---|---|---|---|
| S2' | E58 | H-bond | 2.68 |
| R3' | D105 | Salt bridge | 3.05 |
| E5' | R59 | Salt bridge | 2.71 |
| E5' | N61 | H-bond | 2.81 |
| R11' | D112 | Salt bridge | 2.71 |
| R11' | Y42 | H-bond | 2.98 |
| E14' | R65 | Salt bridge | 2.80 |
| R3' | F110 | Cation-Pi interaction[18] | |

[a] ALFA peptide residues are marked by an apostrophe.
